# Vision-related convergent gene losses reveal *SERPINE3*’s unknown role in the eye

**DOI:** 10.1101/2022.02.25.481972

**Authors:** Henrike Indrischek, Juliane Hammer, Anja Machate, Nikolai Hecker, Bogdan M. Kirilenko, Juliana G. Roscito, Stefan Hans, Caren Norden, Michael Brand, Michael Hiller

## Abstract

Despite decades of research, knowledge about the genes that are important for development and function of the mammalian eye and are involved in human eye disorders remains incomplete. During mammalian evolution, mammals that naturally exhibit poor vision or regressive eye phenotypes have independently lost many eye-related genes. This provides an opportunity to predict novel eye-related genes based on specific evolutionary gene loss signatures. Building on these observations, we performed a genome-wide screen across 49 mammals for functionally uncharacterized genes that are preferentially lost in species exhibiting lower visual acuity values. The screen uncovered several genes, including *SERPINE3*, a putative serine proteinase inhibitor. A detailed investigation of 381 additional mammals revealed that *SERPINE3* is independently lost in 18 lineages that typically do not primarily rely on vision, predicting a vision-related function for this gene. To test this, we show that *SERPINE3* has the highest expression in eyes of zebrafish and mouse. In the zebrafish retina, *serpine3* is expressed in Mueller glia cells, a cell type essential for survival and maintenance of the retina. A CRISPR-mediated knockout of *serpine3* in zebrafish resulted in alterations in eye shape and defects in retinal layering. Furthermore, two human polymorphisms that are in linkage with *SERPINE3* are associated with eye-related traits. Together, these results suggest that *SERPINE3* has a role in vertebrate eyes. More generally, by integrating comparative genomics with experiments in model organisms, we show that screens for specific phenotype-associated gene signatures can predict functions of uncharacterized genes.

## Introduction

Disorders affecting eyes range from subtle vision impairment to blindness and are among the most prevalent diseases in the human population (1). For example, an estimated 76 million people worldwide suffer from glaucoma (2), a disease involving optic nerve damage, and about 1.8 million people are blind due to age-related macular degeneration, a degenerative disease of the central retina affecting retinal pigment epithelium (RPE) and photoreceptor cells (3, 4).

Extensive research in the past decades identified many protein-coding genes with crucial roles in development and maintenance of different tissues and cell types in the eye as well as numerous genes that are associated with genetic eye disorders (1, 5). For example, the RetNet database (6) lists 271 genes associated with heritable retinal diseases. Even though the eye probably represents one of the best studied organs, our knowledge of the genes underlying eye diseases and disorders is still incomplete. For example, linkage analysis in patients with cataract, microcornea, microphthalmia, and iris coloboma identified new genomic loci linked to the diseases; however, the disease-causing genes have remained elusive (7–9). Similarly, the Cat-Map database (10) lists several additional cataract-associated loci where the underlying disease-causing gene has not been identified. Furthermore, there are still thousands of genes that have not been experimentally investigated in detail, leaving many genes where potential eye-related functions remain to be discovered. Indeed, systematic knockouts of 4364 genes in mouse detected ocular phenotypes for 347 genes, with 75% of them not been known as eye-related before (5). This indicates that vision-related genes as well as genes associated with genetic eye disorders remain to be identified and characterized.

Interestingly, many genes that are linked to human eye diseases are inactivated (lost) in non-human mammals that naturally exhibit poor vision (11, 12). For example, subterranean mammals, such as the blind mole rat, naked mole rat, star-nosed mole and cape golden mole, exhibit gene-inactivating mutations in genes implicated in cataract, retinitis pigmentosa, color or night blindness, or macular degeneration in human (e.g. *ABCA4*, *BEST1*, *CRYBA1*, *EYS*, *GJA8*, *GNAT2*, *PDE6C*, *ROM1* and *SLC24A1*) (12–18). Symptoms that characterize these human eye diseases resemble traits found in these subterranean mammals, such as highly-degenerated retinas and lenses and sometimes blindness. Similarly, losses of the short wave sensitive opsin (*OPN1SW*), which is linked to color blindness in humans, occurred in several mammalian lineages such as cetaceans, bats, sloths and armadillos that are consequently inferred to have monochromatic vision (11, 19–23). In addition to losses of vision-related genes, subterranean mammals also exhibit widespread sequence and transcription factor binding site divergence in eye-related regulatory elements (24–26). Such mutations can cause mis-expression of target genes in ocular tissues such as the lens (27). Loss and divergence of vision-related genes and regulatory elements in these mammals is likely caused by the lack of natural selection on maintaining functional eyes in a dark environment. Taken together, previous studies established a clear association between naturally occurring poor vision phenotypes and regressive evolution of the genetic machinery required for functional eyes (11, 17, 24, 25).

Here, we performed a genome-wide screen for genes preferentially lost in independent mammalian lineages with a low visual acuity with the goal of revealing currently uncharacterized genes, where vision-related gene loss patterns would predict vision-related functions. In addition to identifying losses of known vision-related genes, our screen revealed previously unknown losses of several functionally uncharacterized genes, among them *SERPINE3*, which is lost in 18 mammalian lineages that often do not use vision as the primary sense. We show that *SERPINE3* is specifically expressed in eye of zebrafish and mouse. Furthermore, by knockout of *serpine3* in zebrafish, we show that the gene is required for maintenance of a proper retinal lamination and overall eye shape, which confirms the predicted vision-related function. Collectively, our results confirm that *SERPINE3* has functions in vertebrate eyes and our discovery-driven study demonstrates how specific evolutionary divergence patterns can reveal novel insights into gene function (28).

## Results

### A genome-wide screen retrieves genes preferentially lost in mammals with low visual acuity

To uncover potentially unknown vision-related genes, we used the Forward Genomics framework (29) to search for associations between convergent gene losses and poor vision phenotypes in mammalian lineages, where poor vision has independently evolved. Vision is a complex multifaceted trait that may not be easily captured with a single variable. In our study, we decided to select mammals with poor vision based on visual acuity values for two reasons. First, visual acuity describes the ability of an animal to resolve static spatial details, which generally reflects how much an animal relies on vision in comparison to other senses (30). Second, visual acuity data is available for 49 placental mammals with sequenced genomes, enabling a comprehensive genomic screen (Tab. S1). Using visual acuity values, we defined two groups. Low-acuity species have low visual acuity values <1 (log_10_(va)<0), which comprises ten species (three echolocating bats, three rodents and four subterranean mammals) representing seven lineages (Fig. 1A). All other species have visual acuity values >1 and comprise the group with higher visual acuity values.

**Figure 1:**
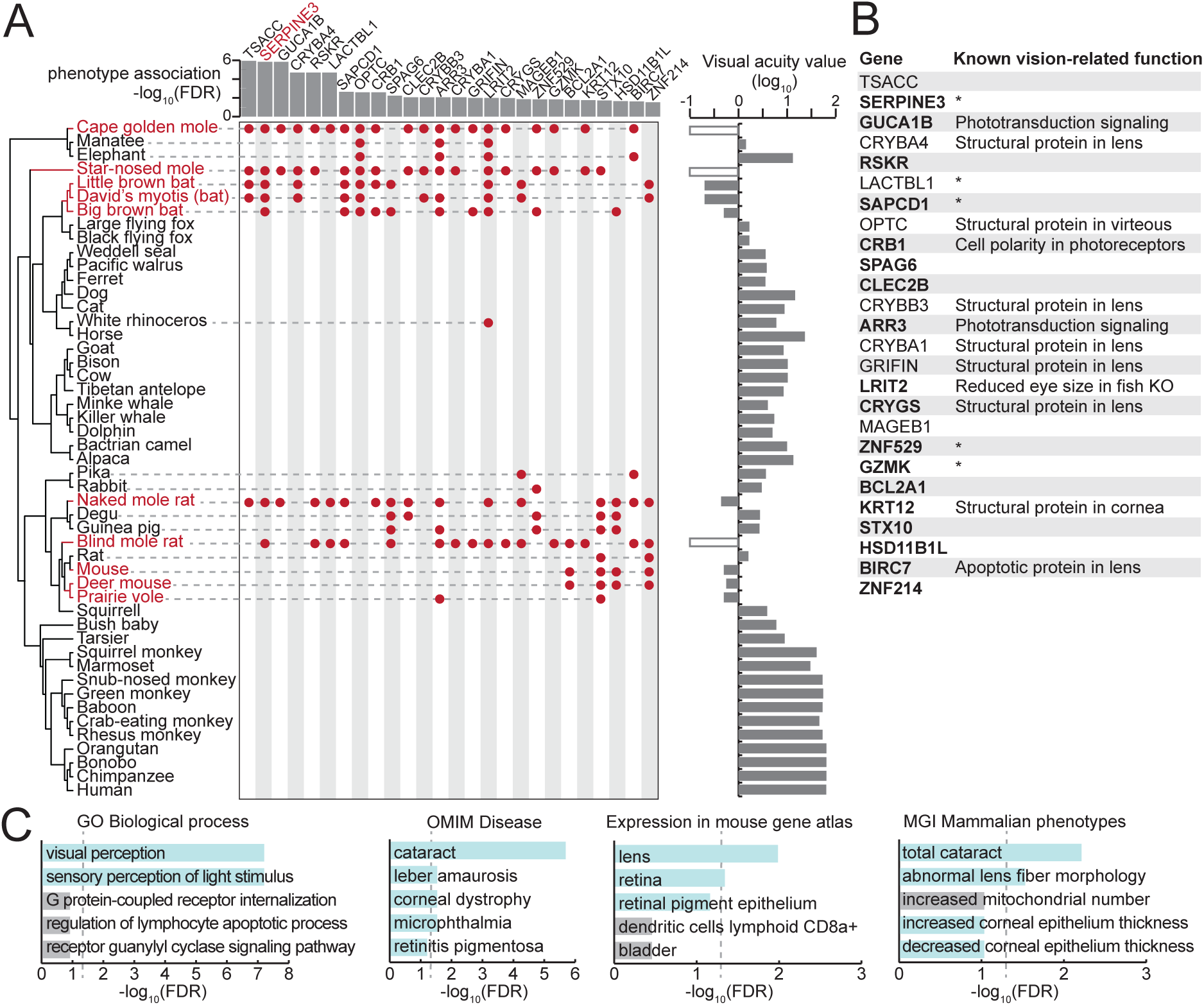
A comparative genomics screen uncovered known and novel vision-related genes. (A) Phylogeny of the species included in our screen (left). Visual acuity values on a log_10_ scale are shown on the right as a bar chart; white boxes indicate the three subterranean mammals that lack acuity measurements but are functionally blind. Low-acuity species (red font) are defined here as species with visual acuity <1 (log_10_(va)<0, results for other thresholds are provided in Tab. S1). At a false discovery rate (FDR) threshold of 0.05, our screen retrieved 26 genes, which are preferentially lost in low-acuity species. Genes lost in individual species are denoted by red dots. The FDR value for the gene loss – phenotype association is shown at the top as a bar chart. (B) List of 26 genes, together with known vision-related functions. Asterisk marks genes that have no known vision-related function but were mentioned in large-scale gene expression data sets of ocular tissues. Genes in bold are expressed in human eyes according to the eyeIntegration database (98) (see Methods). KO - knockout. (C) Functional enrichments of the 26 genes reveals vision-related functions (Gene Ontology, GO of biological processes), associations with human eye disorders (OMIM), expression in ocular tissues in the mouse gene atlas (80) and eye phenotypes in mouse gene KOs (MGI Mammalian Phenotype level 4) among the top five most significant terms. Vision-related terms are shown as blue bars. The dashed line indicates statistical significance in a one-sided Fisher’s exact test after correcting for multiple testing using the Benjamini-Hochberg procedure (FDR 0.05).

Using this classification, we performed a genome-wide screen for genes that exhibit inactivating mutations (frame-shifting insertions or deletions, premature stop codons, splice site mutations, exon or gene deletions) preferentially in low-acuity species, similar to previous screens (31–33). Using a false discovery rate (FDR) cutoff of 0.05, we obtained 26 genes which were each convergently lost in at least three lineages of low-acuity mammals (Fig. 1B, Tab. S2). These genes include several known vision-related genes, such as components of the photoreceptor signal transduction cascade (*GUCA1B*, *ARR3*), a factor required for retinal organization (*CRB1*), lens crystallins (*CRYBA1*, *CRYBB3*, *CRYGS*) and the cornea specific keratin 12 (*KRT12*). The set of 26 genes is enriched for vision-related functions such as the Gene Ontology term “visual perception” (p=5.9e-8), expression in the mouse lens (p=1.0e-2), and human eye diseases (cataract: p=6.0e-3) (Fig. 1C, Tab. S3). This shows that our genome-wide screen successfully retrieved known vision-related genes.

Interestingly, while 14 of the 26 top-ranked hits have no studied role in the eye (Tab. S2), many of these genes are expressed in human eye and *RSKR*, *LACTBL1* and *ZNF529* even cluster in their expression pattern with other retina phototransduction and visual perception genes (34). The preferential loss of these genes in species that exhibit lower visual acuity values and that have convergently lost other known vision-related genes predicts an uncharacterized vision-related function for some of these genes (Fig. 1A).

### *SERPINE3* is convergently lost in low-acuity mammals

We sought to experimentally test this prediction for a gene that is ranked highly in our screen. We focused on the second-ranked candidate, *SERPINE3* (serpin family E member 3), since the first-ranked gene in our screen, *TSACC* (TSSK6 activating cochaperone), encodes a chaperone that is specifically expressed in testis (34, 35) and is therefore unlikely to have an eye-related function. *SERPINE3* is largely uncharacterized and classified as lost in 7 of the 10 low-acuity mammals (Fig. 1, Tab. S2). *SERPINE3* independently accumulated inactivating mutations in all four subterranean species (cape golden mole, star-nosed mole, naked mole rat, blind mole rat). Within bats, *SERPINE3* is inactivated in the two *Myotis* bats and the big brown bat (Yangochiroptera). These three species use echolocation instead of vision as the primary sense for hunting. In contrast, non-echolocating flying foxes (Pteropodidae) that rely more on vision possess an intact *SERPINE3* gene. Finally, varying the threshold used to classify species as having low-acuity vision, consistently retrieves *SERPINE3* as one of the top-ranked hits (Tab. S2), showing that this association is robust to the selected thresholds.

### *SERPINE3* became dispensable in many mammals that do not primarily rely on vision

To explore the evolution of *SERPINE3* in additional mammalian genomes, we made use of an orthology data set generated by the TOGA (Tool to infer Orthologs from Genome Alignments) method (36) that includes 381 additional placental mammalian species that were not part of the genomic screen. Interestingly, this substantially extended data set revealed a number of additional losses of *SERPINE3*, typically in species that do not rely on vision as their primary sense (Fig. 2A). A detailed analysis of inactivating mutations indicates that *SERPINE3* is inactivated in 70 of the 430 analyzed placental mammals and that the gene is convergently lost at least 18 times in placental mammal evolution (Figs. S1-7, Tab. S4).

**Figure 2:**
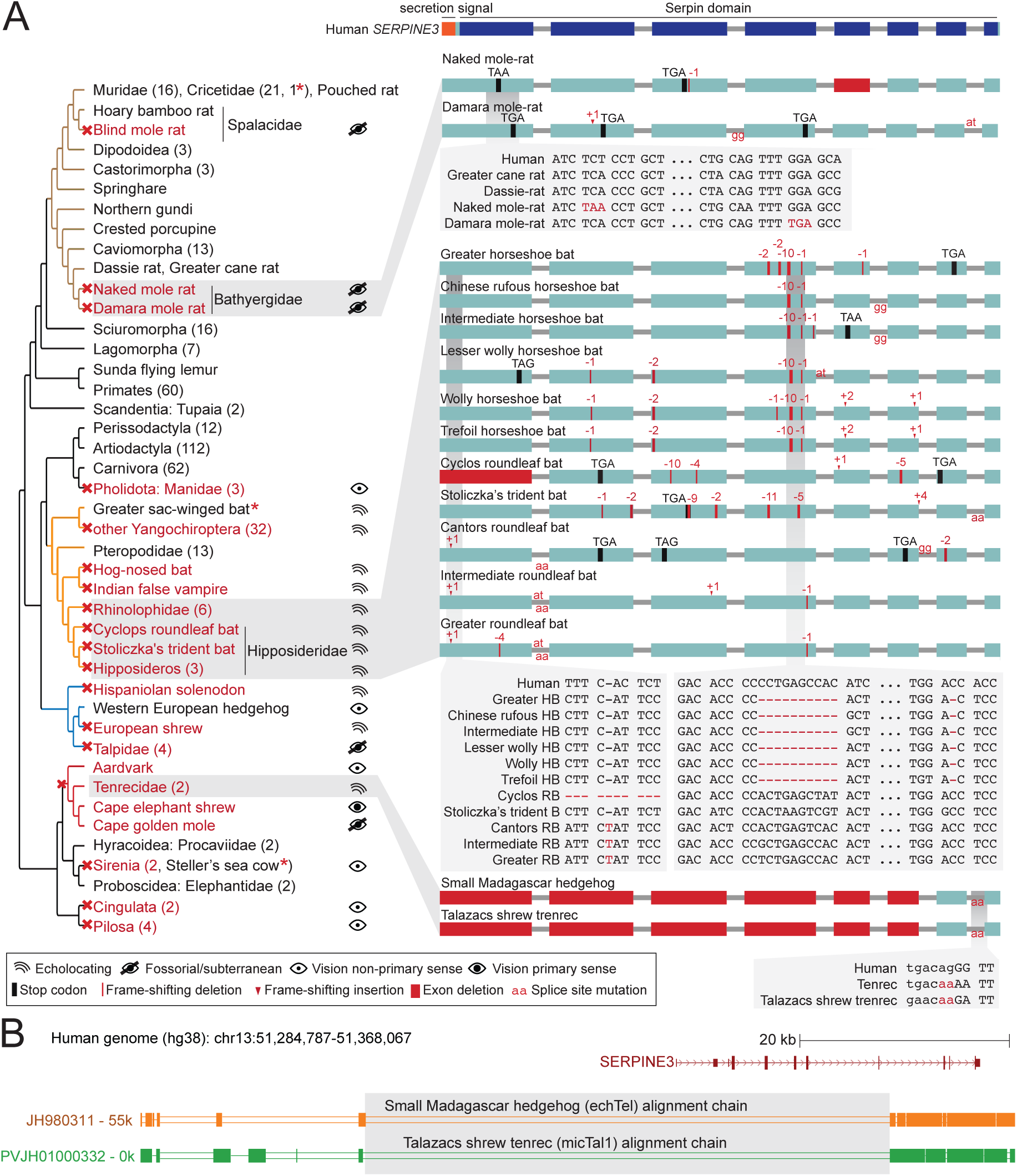
*SERPINE3* gene loss pattern across 430 mammalian species. (A) Left: Phylogeny of mammalian species investigated for the loss of *SERPINE3* with mapped gene loss events indicated as red crosses. Branches of major clades are colored (Rodentia – brown, Chiroptera – orange, Eulipotyphla – blue, Afroinsectiphilia - red). The number of species investigated per clade is specified in parenthesis. For all loss lineages (red font), visual capability (classified as echolocating, fossorial/subterranean, vision as non-primary and primary sense) is displayed as pictograms at the right. Asterisk marks indicate species, where *SERPINE3* evolved under relaxed selection but did not accumulate inactivating mutations. Right: The Serpin protein domain (Pfam) spans all eight protein-coding exons (boxes) of the intact human *SERPINE3* gene (top). Gene-inactivating mutations are illustrated for three clades, with stop codon mutations shown in black, frame-shifting insertions and deletions shown in red and mutated splice site dinucleotides shown between exons in red. Deleted exons are shown as red boxes. Insets show codon alignments with inactivating mutations in red font. RB - roundleaf bat, HB - horseshoe bat. See Fig. S1-7 for detailed plots for all species with gene-inactivating mutations. (B) UCSC genome browser (106) view of the human hg38 assembly showing the *SERPINE3* locus and the orthologous alignment chains of two tenrec species (blocks represent aligning sequence, double lines represent unaligning sequence). A deletion removed the first five protein-coding exons of *SERPINE3* in both species. Shared breakpoints (gray box) indicate that the deletion likely represents an ancestral event in Tenrecidae.

For example, *SERPINE3* is lost in several mammalian clades with nocturnal representatives that have a partially burrowing lifestyle such as pangolins (Manidae), aardvark and armadillos (Cingulata), which are characterized by proportionally small eyes (37, 38) (Fig. 2A, Figs. S1-2). All representatives of these clades use smell and hearing as primary senses for perception of environmental clues and have a poor sense of vision (39). *SERPINE3* has also been lost in the stem lineage of Pilosa (sloths and anteaters). Whereas extant sloths have rather high visual acuity values (10.2 for southern two-toed sloth) (40), the Pilosa ancestor was likely a digging/burrowing species (41), indicating that *SERPINE3* loss originally occurred in a burrowing species. *SERPINE3* is also lost in solenodon, European shrew and tenrecs (Tenrecidae), species with small eyes that use echolocation for close-range spatial orientation (42–44) (Fig. 2B, Figs. S1, S3). For bats, our extended data set shows that *SERPINE3* is convergently lost in seven echolocating lineages, but remains intact in all 13 analyzed Pteropodid bats, revealing a clear pattern of convergent losses of this gene restricted to bat lineages relying on laryngeal echolocation (Figs. S4-6). The sac-winged bat is the only laryngeal echolocating bat with an intact *SERPINE3*; however, selection rate analysis indicates that the gene evolves under relaxed selection (Tab. S5). The available genomes support shared gene-inactivating mutations among many bat species, which likely represent ancestral events (Fig. 2A right). Expanding upon the two convergent losses of *SERPINE3* in subterranean rodents detected in the initial screen, the gene is also convergently lost in the fossorial Damara mole rat and evolves under relaxed selection in the Transcaucasian mole vole, while being intact in all other 63 rodents (Tab. S5, Fig. S7). In the expanded data set, we further uncovered the loss of *SERPINE3* in another clade of fossorial moles (Talpidae) within the order of Eulipotyphla (Fig. S3). A splice site mutation shared between dugong and manatee as well as patterns of relaxed selection indicate an ancestral loss of *SERPINE3* in the ancestor of sirenians, which mostly use their tactile sense for navigation through murky water (45) (Fig. S1, Tab. S5). Consistent with our definition of poor vision based on visual acuity, the Florida manatee has a reduced ability to resolve spatial detail (va = 1.6 (46)) compared to its closest relatives, the African elephant (va = 13.16 (47)), which has an intact *SERPINE3* gene. Finally, *SERPINE3* is also lost in the cape elephant shrew, which is a mostly diurnal species with relatively large eyes (48); however, loss of *SERPINE3* already occurred in the ancestral Afroinsectiphilia lineage (Fig. S1), which was presumably nocturnal (49).

Together, many independent gene losses in species that do not rely on vision as their primary sense predicts a vision-related function for *SERPINE3* that became dispensable in these mammals, leading to convergent *SERPINE3* losses due to relaxed selection.

### *SERPINE3* encodes a putative secreted proteinase inhibitor

Based on sequence homology, *SERPINE3* is classified as a member of the **ser**ine **p**roteinase **in**hibitor (SERPIN) family. Many members of this family are secreted into the extracellular space and inhibit their substrates by covalent binding (50, 51). We performed a SERPINE3 sequence analysis, which revealed that key sequence features of inhibitory serpins are well conserved among placental mammals, suggesting that SERPINE3 also functions as a secreted serine proteinase inhibitor (SI Text, Figs. S8-9).

Serpins have roles in coagulation, angiogenesis, neuroprotection and inflammation, and several serpins have been implicated in human diseases (51, 52). However, the functional role of *SERPINE3* is largely unknown as the gene has never been studied in an animal or cellular model. A thorough literature search revealed that *SERPINE3* is listed (often in the supplement) among many other genes, as differentially expressed in large-scale expression analyses. For example, *Serpine3* is upregulated in the mouse retina in response to overexpressing neuroprotective factors (53). *Serpine3* was also upregulated in complement component 3 (*C3*) knockout mice, which represent a model of the aged retina (54), and downregulated in the eye of knockout mice for *PCARE*, a causal gene for retinitis pigmentosa (55). Together, while this gene was never studied in greater detail, literature clues and the striking convergent gene loss pattern in mammals with poor vision suggest that *SERPINE3* may have a functional role in the eye.

### *Serpine3* is specifically expressed in Müller glia in the adult zebrafish retina

To test the prediction that *SERPINE3* has an eye-related function, we first analyzed its expression pattern in adult zebrafish (Fig. 3). Zebrafish has proven as valuable model species for the study of eye genetics as it has a cone-dominated retina (∼60% cones, ∼40% rods) (56), similar to the central human retina. Furthermore, zebrafish have a single *serpine3* ortholog that is located in a context of conserved gene order (Fig. 4A), which makes zebrafish a suitable model for investigation of *serpine3* expression and function. Reverse transcription quantitative PCR (RT-qPCR) analysis of biological triplicates revealed that the highest *serpine3* expression is in the eye, followed by significantly lower expression in the brain (one-sided t-test, p=0.031, Fig. 3A). *Serpine3* expression was not detectable in the other tested adult tissues (heart, intestine, liver and testis).

**Figure 3:**
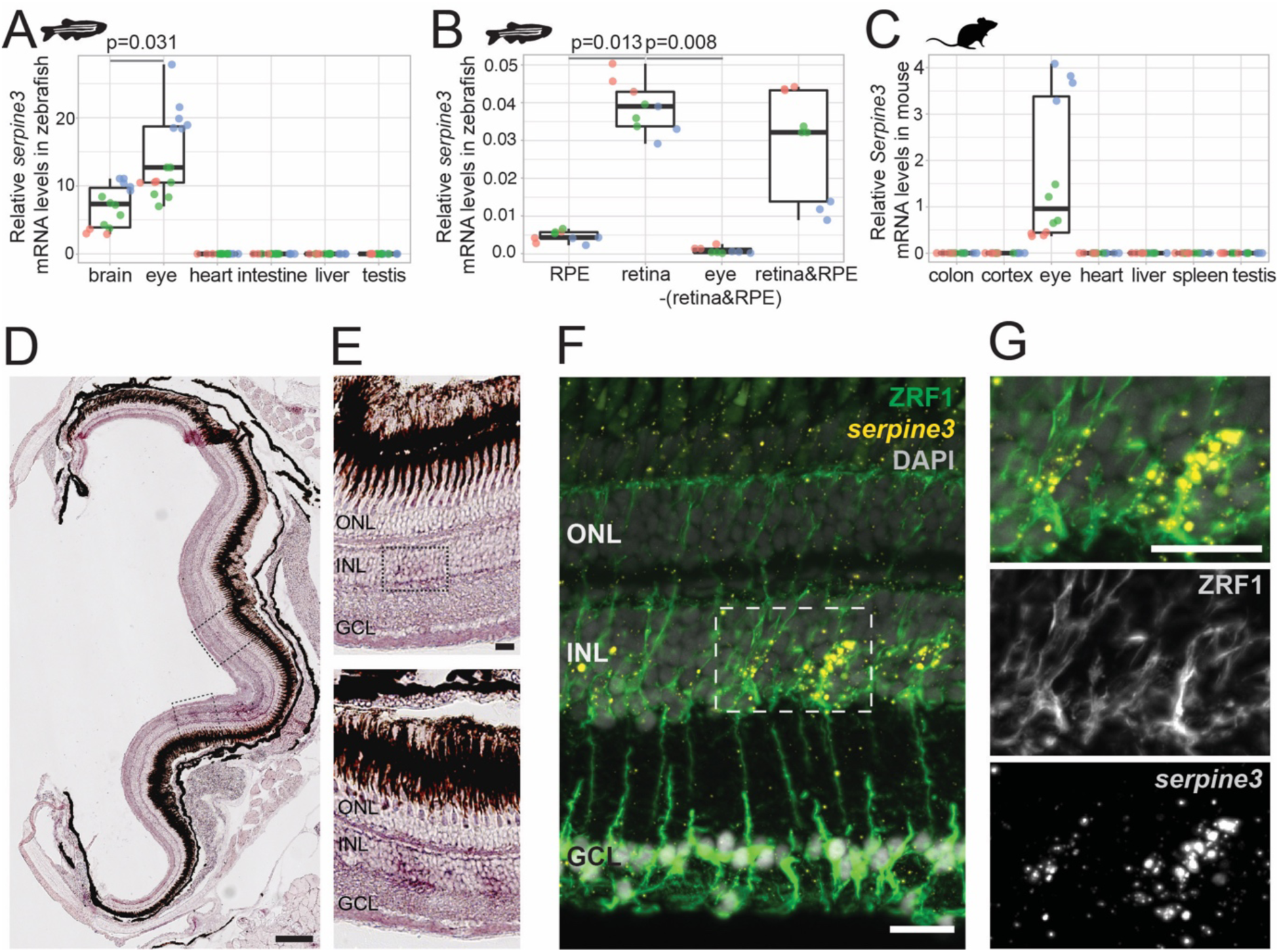
*Serpine3* is expressed in zebrafish and mouse eyes. (A) Expression of zebrafish *serpine3* mRNA in relation to the reference gene *rpl13a* measured with RT-qPCR. *Serpine3* is expressed in brain and eye but not in heart, intestine, liver or testis of adult zebrafish. Expression was significantly higher in eye compared to brain (one-sided t-test). (B) *Serpine3* mRNA expression in different tissues of the zebrafish eye in relation to the reference gene *actb* measured with RT-qPCR. *Serpine3* is specifically expressed in the retina but not in other tissues of the eye. The expression level is significantly higher in retina without RPE compared to RPE only (two-sided t-test). (C) *Serpine3* mRNA expression in mouse in relation to the reference gene *Rpl27* measured with RT-qPCR. *Serpine3* is specifically expressed in the eye but not in colon, cortex, heart, liver, spleen and testis. Technical replicates are shown in the same color. (D-G) *Serpine3* mRNA expression pattern in zebrafish retina. Chromogenic *in situ* hybridization (ISH, D-E) shows localized expression of *serpine3* (purple) in the retina, specifically in the inner nuclear layer (inlet). (F-G) Fluorescence *in situ* hybridization shows that *serpine3* mRNA expression (yellow) is localized to cell bodies of Mueller glia cells. Filaments of Mueller glia cells are marked by the glial fibrillary acidic protein (ZRF1 antibody, green). Nuclei are stained with DAPI (white). Scale bar is 200 µm in (D) and (F) and 20 µm in (E) and (G). INL – inner nuclear layer, ONL – outer nuclear layer, GCL – ganglion cell layer.

**Figure 4:**
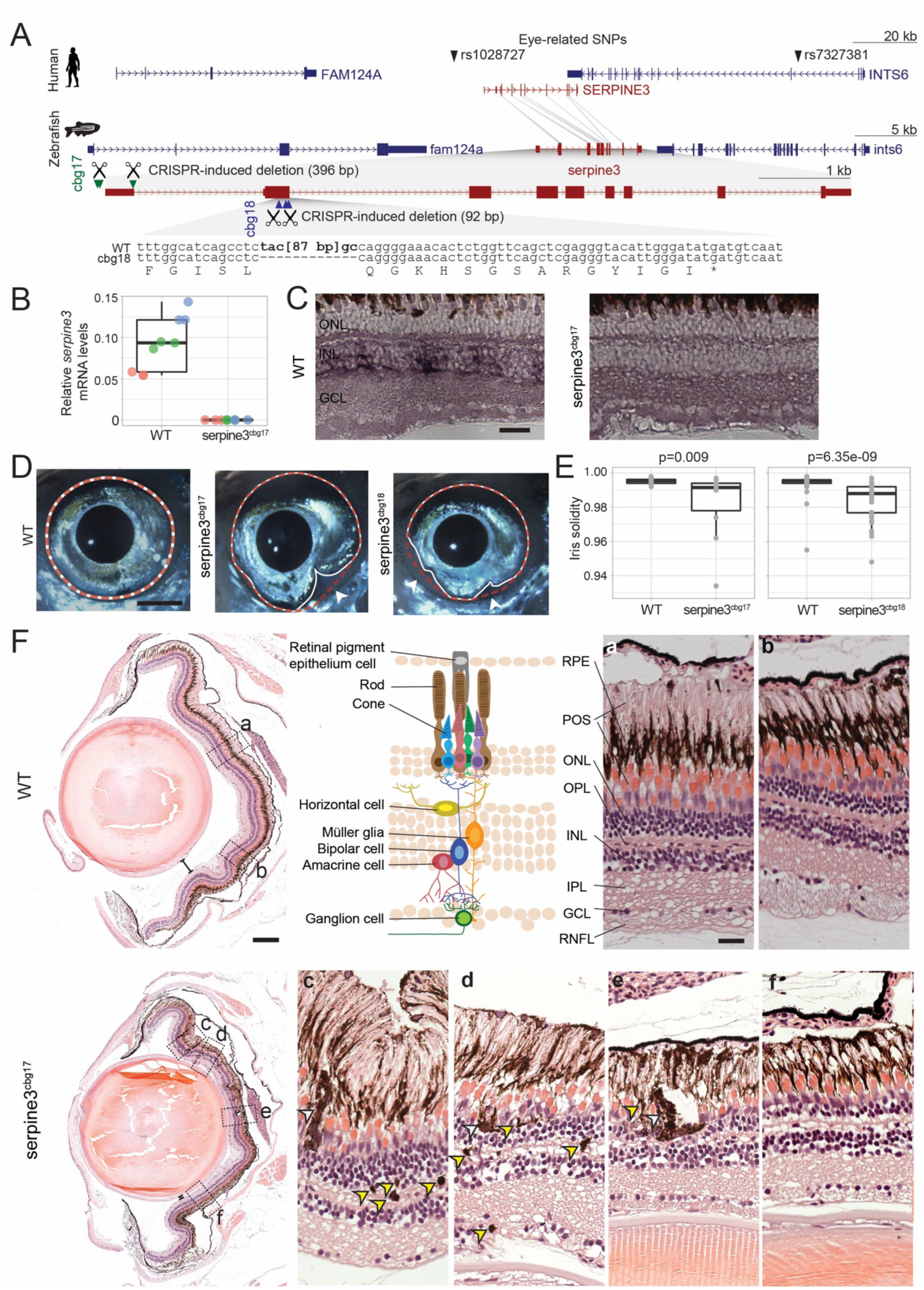
*Serpine3* knockout in zebrafish causes defects in eye shape and retinal layering. (A) UCSC genome browser visualization of the *SERPINE3* genomic locus in human (hg38 assembly, top) and zebrafish (danRer11 assembly, bottom) shows that both species have a 1:1 ortholog with the same number of coding exons in a conserved gene order context. In the human locus, two single nucleotide polymorphisms (SNPs) are in linkage with *SERPINE3* and associated with eye phenotypes. In zebrafish, we used CRISPR-Cas9 to generate two independent knockout (KO) lines. The position of guide RNAs is indicated as scissors. In the *serpine3*^cbg17^ line, we deleted the promoter and first exon. In the *serpine3*^cbg18^ line, we introduced a 92 bp frame-shifting deletion in exon 2 (coding exon 1) that results in three early stop codons in the original reading frame. (B) Relative expression of *serpine3* mRNA in wild type (WT) zebrafish and *serpine3*^cbg17^ individuals (n=3 each) quantified by RT-qPCR relative to the expression of *rpl28*. *Serpine3*^cbg17^ fish do not express *serpine3* mRNA. Technical replicates are shown as individual data points, the same color encoding one biological replicate. (C) *In situ* hybridization showing that *serpine3* is expressed in the inner nuclear layer (INL) of WT zebrafish but not in the homozygous *serpine3*^cbg17^. Scale bar = 25 µm. (D) *Serpine3* knockout leads to changes in eye shape in adult, homozygous knockout (KO) fish of *serpine3*^cbg17^ and *serpine3*^cbg18^ lines in comparison to their WT siblings (18 and 11 months, respectively). In WT, the eye shape almost perfectly corresponds to the concave shape of the iris (overlay of white and red dotted lines). In contrast, many KO individuals have alterations in eye shape, evident by notches in white line that follows the iris. Scale bar = 1 mm. (E) Iris solidity (ratio of eye shape/ concave eye shape) significantly differs between WT and KO siblings for both the *serpine3*^cbg17^ (16 vs. 10 eyes) and the *serpine3*^cbg18^ (40 vs. 40 eyes) line. A Wilcoxon Rank sum test was used. (F) Hematoxylin/eosin histology staining of the eye of *serpine3*^cbg17^ fish (22 months) reveals histological differences in comparison to their WT siblings (dorsal top, ventral bottom). In comparison to WT, distance between lens and retina of *serpine3*^cbg17^ fish is reduced (distance bars). The WT retina (top) has a distinct lamination with clear separation of the single retinal layers (a, b) as shown in the schematic (RPE – retinal pigment epithelium layer, POS – photoreceptor outer segment, ONL – outer nuclear layer, OPL – outer plexiform layer, INL – inner nuclear layer, IPL – inner plexiform layer, GCL – ganglion cell layer, RNFL – retinal nerve fiber layer). Although all retinal layers are present in *serpine3*^cbg17^ fish, the layering appears distorted and the density of cells is reduced (c-f). Specifically, the RPE cells display an altered distribution and even local clusters (empty arrows), and displaced pigmented cells emerge in all retinal layers (yellow arrows). Scale bar in the overviews = 200 µm, scale bar in the magnifications = 20 µm.

To better characterize *serpine3* expression in the eye, we next performed RT-qPCR on samples of RPE, retina and eye without retina and RPE (Fig. 3B). *Serpine3* is expressed significantly higher in the retina compared to RPE only (two-sided t-test, p=0.013, Fig. 3B). It is barely detectable in whole eye without retina and RPE (two-sided t-test, eye vs. retina, p=0.008, Fig. 3B), suggesting that the expression signal obtained from whole eye RT-qPCR mostly originates from expression in retina.

To specify in which cell types *serpine3* is expressed, we finally performed *in situ* hybridization (ISH) on adult zebrafish retinas. *Serpine3* expression is detected in the inner nuclear layer throughout the retina with stronger signals in the ventral region close to the optic nerve (Fig. 3D, E). Co-staining with different cell type specific markers reveals expression of *serpine3* in a fraction of glial fibrillary acidic protein (*gfap*)-positive Mueller glia (MG) cells (Fig. 3F, G, Fig. S10), whereas expression of *serpine3* mRNA was not detected in bipolar or amacrine cells (Fig. S11).

To also explore *SERPINE3* expression in mammals, we first used RT-qPCR to analyze *Serpine3* expression in mouse. Similar to zebrafish, mouse *Serpine3* expression is highest in whole eye, whereas expression was barely detectable in colon, cortex, heart, liver, spleen and testis (Fig. 3C). Next, we analyzed *SERPINE3* expression in other mammals and vertebrates using publicly available expression data sets. Despite sparser data, we found evidence for expression of *SERPINE3* in eyes of human, rat, cat, cow and chicken, which suggests a conserved expression pattern and a role in vertebrate vision (Tab. S6).

In summary, our experiments in zebrafish and mouse together with available data sets of different vertebrates show that *SERPINE3* is expressed in the vertebrate eye, specifically in zebrafish in MG cells.

### Knockout of *serpine3* in zebrafish leads to morphological defects in the eye including the retina

Next, we tested whether *serpine3* inactivation results in an eye phenotype. To this end, we deleted the transcription start site of *serpine3* with CRISPR-Cas9, denoted as the *serpine3*^cbg17^ allele (Fig. 4A, Figs. S12-13). Adult homozygous *serpine3*^cbg17^ fish were viable and fertile. Using RT-qPCR and ISH, we confirmed that the deletion of the transcription start site completely abolished *serpine3* expression in retinae of adult homozygotes for *serpine3*^cbg17^ compared to wild type (WT) siblings (Fig. 4B, C).

We found that adult *serpine3*^cbg17^ individuals frequently showed notches in the iris of one of the eyes (4 of 5 individuals, Tab. S7, white arrow in Fig. 4D), which affected the eye’s overall shape. To confirm that these notch-caused shape deviations are caused by inactivation of *serpine3*, we generated an independent line, where we introduced an early frameshift in *serpine3’s* coding exon 1 with CRISPR-Cas9, denoted as the *serpine3*^cbg18^ allele (Fig. 4A, Figs. S12-13). *Serpine3*^cbg18^ fish also showed a high frequency of notches in 13 of 20 individuals (Tab. S7). To quantify this shape deviation, we calculated iris solidity, which compares the area ratio of the eye’s outer shape (white line) and its concave shape (red dotted line in Fig. 4D). Eyes of both *serpine3*^cbg17^ and *serpine3*^cbg18^ individuals had a significantly reduced solidity in comparison to their WT siblings (Wilcoxon rank sum test, p=0.009 and p=6.35e-9, respectively, Fig. 4E). Iris circularity, another descriptor of the eye shape deviation, is also significantly reduced in homozygous individuals of both lines (Fig. S14). Together this indicates that mutations in *serpine3* cause eye shape deviations in zebrafish.

We next investigated the ocular morphology of adult fish of both lines and their WT siblings using hematoxylin/eosin sections (Fig. 4F; Figs. S15-16). While ocular structure and optic nerves of all inspected eyes are largely normal (Fig. S15), the distance between retina and lens is reduced in *serpine3*^cbg17/18^ KO fish (Fig. 4F, Fig. S16). A detailed inspection of the retinal organization revealed that although all retinal layers were present, the mutant retinae are generally less organized and structured compared to WT siblings. Retinal cells appear reduced in number and less densely packed. Most prominently, we noticed that rod outer segments and the pigmented RPE cells, which were aligned in WT fish, were not clearly separated in *serpine3*^cbg17/18^ fish (Fig. 4F c-e, Fig. S16). Furthermore, we observed large clusters of pigmented cells in the photoreceptor layer (empty arrows, Fig. 4F c-e) as well as single displaced pigmented cells in all retinal layers (yellow arrows, Fig. 4F c-e). Similar alterations in retinal structure were detected in *serpine3*^cbg18^ fish (Fig. S16). This shows that KO of *serpine3* in zebrafish results in morphological defects in the eye, characterized by differences in eye shape and retinal organization.

### Polymorphisms near human *SERPINE3* are associated with human eye phenotypes

We next analyzed recently published Genome Wide Association Study data for human single nucleotide polymorphisms (SNPs) associated with eye-related traits. This analysis revealed two such SNPs that are in linkage disequilibrium with *SERPINE3* (Fig. 4A, Tab. S8). rs1028727 is located ∼10 kb upstream of the *SERPINE3* transcription start site and is associated with a decreased area of the optic nerve head (57). rs7327381 is located ∼97 kb downstream of the *SERPINE3* transcription start site and is associated with an increase in corneal curvature (58). This suggests that human SNPs linked to *SERPINE3* are associated with eye phenotypes, supporting a putative ocular function of human *SERPINE3*.

## Discussion

Our study combines comparative genomics to predict genes having vision-related functions and experiments in zebrafish to confirm this prediction for a top-ranked candidate, the uncharacterized *SERPINE3* gene. By conducting a genome-wide screen for genes that are preferentially lost in mammals with low visual acuity values, we uncovered both known vision-related genes as well as several genes that have no known eye-related function. One of the top-ranked candidates is *SERPINE3*, which we found to be independently lost at least 18 times in mammalian evolution, preferentially in species that do not use vision as the primary sense. For mouse and zebrafish, we show that the highest *SERPINE3* expression is in the eye, which is corroborated by available expression data of other vertebrates. By generating the zebrafish *serpine3*^cbg17^ and *serpine3*^cbg18^ knockout (KO) lines, we show that inactivation of this gene results in abnormal eye shape and retinal lamination, revealing an eye-related function for *serpine3*. This is further supported by *SERPINE3*-linked polymorphisms that are associated with eye-related traits in human.

In zebrafish, *serpine3* is expressed in a fraction of Mueller glia (MG) cells. MG are the major retinal macroglia and perform numerous functions. By removing waste products and secreting (neuro)trophic substances and signaling molecules, they maintain the blood-retinal barrier and regulate vascularization (59). Most importantly, MGs are essential for the long-term viability of photoreceptors and other neuronal cell types (60). Our *serpine3* KO fish exhibit a disorganization of RPE cells, a phenotype that resembles those previously observed in experiments that perturb MG function. For example, selective ablation of MG in adult mice results in eye defects including aggregation of RPE cells and displacement of pigment granules in the ganglion cell layer (61). Interestingly, we also observed that *serpine3* KO affects eye shape, which cannot be readily explained by a direct MG-mediated effect. However, MG span the entire retina, connecting the extracellular space of retinal neurons, the vitreous and the capillaries at the apical retina (59, 62). It is therefore possible that a secreted protein, as predicted for serpine3 based on its conserved signal peptide, may also affect other eye subtissues. Finally, in zebrafish, MG are able to regenerate retinal neurons upon injury. However, *serpine3* does not seem to be involved in this process as during stress-induced regeneration, it is upregulated in resting MG that do not proliferate (63).

In mammals, the *SERPINE3* gene loss pattern and expression profile in several species also support an eye-related function; however, the cell type expression pattern may differ between mammals and zebrafish. While our experiments in zebrafish show *serpine3* expression in MG, which is in agreement with (63), in mouse and human, *SERPINE3* seems to be expressed in RPE (63–65) indicating that cell type specificity may differ between mammals and fish. As *SERPINE3* is likely a secreted extracellular protein, it is possible that it has a similar function in mammals and zebrafish, despite secretion from different cell types. Whether this is the case or whether *SERPINE3* function and protein expression pattern in the mammalian eye differs remains to be explored in future studies. At the molecular level, this question may be addressed by investigating whether the molecular targets of *SERPINE3* are conserved among vertebrates.

Of particular interest is elucidating the functional role of *SERPINE3* in human. Several pieces of evidence indicate a potential role of this gene in anti-inflammatory processes and retinal survival. *SERPINE3* is upregulated in human patients with age-related macular degeneration (66), a progressive eye disease that is linked to chronic inflammation and wound healing. Consistent with a role in retinal survival, mouse *Serpine3* is upregulated after experimental overexpression of neurotrophin-4, a neuroprotective factor that promotes retinal survival (53). Furthermore, *SERPINE3* is a hallmark gene of differentiated, healthy human RPE cells (67–69) that also have neurotrophic functions in the retina. It is thus conceivable that perturbation of proper *SERPINE3* expression or function may influence age-related diseases or human eye phenotypes, as indicated by human polymorphisms that are linked to *SERPINE3*.

In addition to *SERPINE3*, our screen also revealed other candidate genes for which unknown eye-related functions are plausible. The *LACTBL1* gene encodes a putative serine proteinase (70) and has an expression pattern similar to retina phototransduction genes (34). Furthermore, 20 kb upstream of *LACTBL1* is a linked SNP (rs10158878) that is associated with refractive error in human (GWAS catalog, (71)). Another uncharacterized candidate gene with a strong visual acuity associated loss pattern is *SAPCD1*. A missense mutation (rs6905572) within this gene is associated with macular degeneration (dbSNP, (72)). Finally, *LRIT2* is linked to SNPs (rs12217769 and rs745480) that are associated with macular thickness and refractive error in human. Whereas this gene was not functionally characterized at the time our screen was conducted, a recent study showed that *lrit2* knockdown led to a reduction in eye size in zebrafish (73). Thus, in addition to *SERPINE3*, for which we performed an initial functional characterization in zebrafish, our screen uncovered other promising, uncharacterized candidates that may have an eye-related function. Overall, this highlights the potential of comparative genomics to shed light on the functional roles of less characterized genes and to help to further identify human disease-causing genes (74).

## Materials and Methods

### Visual acuity values

We used publicly available visual acuity measurements (40, 75) (Tab. S1 lists all primary references) to classify placental mammals into visual high-acuity and low-acuity species. Visual acuity can be measured by either behavioral experiments or calculated from the eye axial diameter, the peak ganglion cell density and a correction factor for diel activity (40). For species for which both measures were available, we used the behavioral visual acuity as this measure is more accurate (40). For species that lack visual acuity data, we used available visual acuity measurements of closely related species of the same genus or family (Tab. S1). Three subterranean mammals lack available visual acuity measurements (cape golden mole, star-nosed mole, blind mole rat) but exhibit highly degenerated eyes. We therefore assumed a visual acuity of zero. In total, visual acuity values were obtained for 49 placental mammals that were included in a previously-generated whole genome alignment (76). Using a visual acuity threshold of one, we considered ten mammals (cape golden mole, naked mole rat, star-nosed mole, blind mole rat, big brown bat, little brown bat, David’s myotis bat, mouse, prairie vole, deer mouse) representing seven lineages, as low-acuity vision species. All other 39 species with visual acuity greater than one were considered as high-acuity vision species.

### Forward genomics screen

To screen for genes that are preferentially lost in low-acuity placental mammals, we used a previously-generated data set of inactivated genes (33). Briefly, gene losses were detected with a pipeline that searches for gene-inactivating mutations and performs a number of filtering steps to distinguish between real mutations and artifacts related to genome assembly or alignment issues and exon-intron structure changes. A genome alignment with human (hg38 assembly) as the reference (76), human genes annotated by Ensembl (version 87) (77) and principal isoforms from the APPRIS database (78) were used as input. Based on the relative positions of inactivating mutations, a value measuring the maximum percent of the reading frame that remains intact (%intact) was computed for each gene and species. A gene was classified as lost if %intact was <60%. A gene was classified as intact if %intact was ≥ 90%.

To search for genes that tend to have lower %intact values in the low-acuity group, we adopted the Forward Genomics approach (16, 33). We excluded genes that had missing data due to assembly gaps for more than 50% of low- or high-acuity species. We used phylogenetic generalized least squares (79) to account for phylogenetic relatedness and ranked genes by the Benjamini-Hochberg corrected p-value (FDR<0.05). We further extracted genes that tend to be conserved in high-acuity species by requiring that a gene was classified as intact in ≥80% and classified as lost in ≤10% of high-acuity species. Finally, to detect convergent gene losses, we required that a gene was lost in species representing at least three of the seven independent low-acuity lineages.

To test whether the identification of *SERPINE3*, which we experimentally investigated, is robust to the selected visual acuity threshold, we re-ran the screen after increasing or decreasing the threshold while keeping other parameters constant. Using a more inclusive definition of low-acuity species by increasing the visual acuity threshold to two, which additionally considered manatee, the two flying foxes and rats as low-acuity species, identified a total of six genes at an FDR of 0.05, with *SERPINE3* at the first rank (Tab. S2). Using a more restrictive definition of low-acuity species by decreasing the visual acuity threshold to 0.5 identified 53 genes at an FDR of 0.05 with *SERPINE3* at rank 7 (Tab. S2).

### Enrichment analysis

Enrichment analysis was performed using the Enrichr web service (80), which uses a two-sided Fisher’s exact test and corrects for multiple testing with the Benjamini-Hochberg method.

### Investigating *SERPINE3* in additional genomes

To explore conservation and loss of *SERPINE3* in additional mammalian genomes that became available since our initial screen, we used the TOGA method (Tool to infer Orthologs from Genome Alignments) (36). TOGA uses pairwise alignments between a reference (here human hg38) and a query genome, infers orthologous loci of a gene with a machine learning approach, and uses CESAR 2.0 (81) to align the exons of the reference gene to the orthologous locus in the query. TOGA then classifies each transcript by determining whether the central 80% of the transcript’s coding sequence encodes an intact reading frame (classified as intact) or exhibits at least one gene-inactivating mutation (classified as potentially lost). If less than 50% of the coding sequence is present in the assembly, the transcript is classified as missing.

Focusing on the evolutionarily conserved human *SERPINE3* Ensembl (version 104) transcript ENST00000524365 (82), we analyzed the TOGA transcript classification for 418 assemblies that were not used in the initial screen (Tab. S4). These assemblies represent 381 new placental mammal species. For species, where TOGA classified the *SERPINE3* transcript as missing, we inspected the orthologous alignment chain to distinguish intact and lost orthologs from truly missing orthologs due to assembly gaps. Since assembly base errors can mimic false gene losses (32, 83), we further analyzed species for which only one or two inactivating mutations were detected. For these species, we require that (i) at least one inactivating mutation is shared with a closely-related species, or (ii) the mutation is also present in a different assembly of the same species. If that was not the case, we validated inactivating mutations with raw sequencing reads by aligning the genomic sequence around the mutation against the NCBI short read archive (SRA queried via NCBI megablast) (84). Intactness of *SERPINE3* remains unclear for four species (Tab. S4). In order to map *SERPINE3* loss events on the phylogenetic tree, we searched for gene-inactivating mutations that are shared among phylogenetically related species, where parsimony indicates that these mutations and thus gene loss likely occurred in their common ancestor. Those mutations are shown in boxes in Fig. S1-7, all other mutations are the output of CESAR.

### Selection analysis

For species with an unclear *SERPINE3* loss status (mole vole, steenbok, okapi, fox, Steller’s sea cow) and species or clades that have an intact *SERPINE3* but many close relatives have lost the gene (greater sac-winged bat, European hedgehog, elephants), we tested whether *SERPINE3* evolves under relaxed selection. To this end, we used RELAX from the HyPhy suite (85) to test whether selection pressure was relaxed (selection intensity parameter K<1) or intensified (K>1) in this species or clade, which we labeled as foreground (Tab. S5). We restricted the analysis to 327 *SERPINE3* that are intact and complete (middle 80% of the coding sequence present) and treated those as background. Codon sequences were obtained from TOGA and aligned with MACSE v2.0 (86). This procedure was repeated using only one foreground species/clade at the time. The species tree used for the analysis is in Data set S2.

### Protein sequence analysis and structure prediction

Signal peptides and the cellular location were predicted with the SignalP 5.0 webserver (87) and DeepLoc 1 (88), respectively, for all intact and complete SERPINE3 protein sequences as defined above. Protein sequences were aligned with muscle (89) and visualized with Jalview (90) (Data set S3).

The three-dimensional structure of human SERPINE3 was retrieved from the AlphaFold2 web server (91) (Data set S4). We calculated the root mean square distance (RMSD) to all homologous chains of existing crystal structures of close serpin relatives in native state (Tab. S9) after structural alignment in PyMOL (92). For each crystal structure, we averaged the RMSD for all chains.

### Mining gene function, expression and genetic variation sources

Information on the function of genes discovered in our screen was obtained from GeneCards database (93), UniProt (94), Ensembl (82), Proteomics DB (95), the Human protein atlas (96), the Expression atlas and Single cell expression atlas (97). Expression in human eye for each candidate gene were obtained by averaging expression over all healthy, primary RNA-Seq data sets per tissue (cornea, RPE, retina) provided by the eyeIntegration database (98). Cell lines were not included in the average. A gene was considered to be expressed if the Transcripts Per Million (TPM) value was >100 in cornea, RPE, or retina. Expression of *SERPINE3* was further assessed by retrieving primary data sets from FantomCat (99), GEO profiles (100), and Bgee (101). Tab. S6 provides the list of all data sets.

Phenotype associations of SNPs located in loci of interest were investigated based on the GWAS catalog (102), dbSNP (103) and PheGenI (104). Linkage of SNPs with a candidate gene in 30 human populations was investigated based on the GWAS catalog. To evaluate possible functional consequences of the respective SNPs, we overlapped their (projected) coordinates with regulatory elements from ENCODE for human (hg38) and mouse (mm10) via the web-based server SCREEN v. 2020-10 (105). Additionally, we investigated eye- and retina-associated regulatory elements in the Ensembl and UCSC genome browsers (82, 106).

### Animal husbandry

Adult zebrafish (*Danio rerio*, AB line) were maintained at 26.5°C with a 10/14 h dark/light cycle (107). Embryos and larvae were raised at 28.5°C in the dark until six days old. For phenotyping, we used adult fish of both *serpine3* KO lines (*serpine3*^cbg17^:19 months, *serpine3*^cbg18^: 11 months) generated in this study as well as their WT siblings. WT mice (*Mus musculus*, C57BL/6JOlaHsd line) were maintained in a barrier system at 20-24°C with a 12/12 h dark/light cycle.

### Expression analysis by RT-qPCR

Adult zebrafish >12 months were sacrificed by rapid cooling after anesthesia with MESAB. Adult mice (2 months, male) were sacrificed by cranial dislocation after carbon dioxide anesthesia. Tissues were dissected in ice-cold phosphate buffer (PBS), frozen in liquid nitrogen and stored at -80°C until further use. RNA was extracted from lysed, homogenized tissue with RNeasy mini or midi kits (Qiagen) according to manufacturer’s instructions and reprecipitated if necessary. The RNA Integrity Number (RIN) was >7 for all tissues except spleen (RIN>6). Intact total RNA was reverse transcribed into cDNA using the ProtoScript^®^ II First Strand cDNA Synthesis Kit (NEB) according to manufacturer’s instructions with random primers. RT-qPCR was performed after addition of SybrGreen (Roche). Expression relative to a normalization gene was calculated from Ct values according to the efficiency and delta delta Ct method. Specifically, relative ratios of *Serpine3*/*serpine3* expression (zebrafish forward: GAGACCCAAAACCTGCCCTT, reverse: AGCCGGAAATGACCGATATTGA, mouse forward: TGGAGCTTTCAGAGGAGGGTA, reverse: GATACTGAAGACAAACCCTGTGC) were obtained by using *rpl13a* (forward: TCTGGAGGAACTGTAAGAGGTATGC, reverse: AGACGCACAATCTTGAGAGCGA) or *actb* (forward: CGAGCAGGAGATGGGAACC, reverse: CAACGGAAACGCTCATTGC) as reference gene for zebrafish and *Rpl27* as reference gene for mouse (forward: TTGAGGAGCGATACAAGACAGG, reverse: CCCAGTCTCTTCCCACACAAA). At least three biological replicates per sample group were analyzed, which were each represented by the average normalized relative ratio of three to six technical replicates.

### ISH and FISH

For ISH and immunostainings, fish were scarified, eyes dissected and fixed in 4% paraformaldehyde/ 0.1 M PBS after removal of the lens. Eyes were embedded in gelatin/sucrose and sectioned (14 µm) with a cryostat. The *serpine3 in situ* probe spans the coding exons 3-8 (transcript: ENSDART00000132915.2). Using primers with restriction enzyme cut sites (forward, Not1: TAAGCA GC GGCCGCGTAAAAGTGCCCATGATGTACCAG, reverse, BamHI: TAAGCA G GATCCACAACTCGACCTATAAACAGCAAC), *serpine3* cDNA was amplified from total cDNA and cloned into the pCRII-topo vector (Invitrogen). The antisense probes were transcribed with SP6 polymerase, using a DIG-labeled NTP mix (Roche diagnostics). The (fluorescent) ISH were conducted as previously described with minor modifications (108): Hybridization and washing steps were performed at 60°C. For chromogenic *in situs*, sections were incubated with anti-digoxigenin-AP (Roche), diluted 1:4000 in DIG-blocking reagent (Roche) at 4°C overnight and subsequently developed with NBT/BCIP (Roche). For FISH, sections were washed in PBS immediately after quenching. Sections were blocked for 1 h with 2% blocking reagent in MABT (Perkin-Elmer) and then incubated with anti-digoxigenin-POD (Roche), diluted 1:500. The signal was detected with the TSA Plus Cy3/Cy5 kit (Perkin-Elmer).

### Immunohistochemistry

Immunostainings for glial fibrillary acidic protein (ZRF1 from DSHB, 1:200), choline O-acetyltransferase (AB144P from Millipore, 1:500) and protein kinase C alpha (SC-208 from Santa-Cruz, 1:500) were performed after completion of the FISH protocol according to (108). For chat, antigens were retrieved in preheated 10 mM sodium citrate buffer for 6’ at 85°C. Sections were washed in PBS and 0.3% PBSTx prior to primary antibody incubation. Following the protocol of (108), we washed the sections three times in PBSTx, and incubated in anti-goat or anti-rabbit IgG (H+L) Alexa 488-conjugated secondary antibodies (Invitrogen, 1:750).

### Generation of KO lines and genotyping

To generate *serpine3* KO lines, deletions were introduced using the CRISPR-Cas9 system. Guides were chosen considering efficiency predictions of the IDT DNA CRISPR-Cas9 guide checker and ChopChopV2 (109) on the zebrafish assembly danRer7 and ordered as Alt-R CRISPR-Cas crRNA from IDT DNA. For each line, three guides were simultaneously injected into one-cell stage zebrafish embryos as ribonucleoprotein delivery using the Alt-R CRISPR-Cas system following the manufacturer’s instructions (0.5 fmol crRNA per embryo per guide, 0.68 ng Cas9 protein per embryo). The expected deletions were confirmed by Sanger sequencing in several founder individuals, one of which was chosen as the founder of each line (Fig. S12). Heterozygous cbg17 and cbg18 zebrafish were further outcrossed to WT fish for several generations and then bred to homozygosity.

More specifically, for s*erpine3*^cbg17^, we abolished *serpine3* transcription by deleting the single transcription start site, which is supported by activating histone marks in zebrafish and is also well conserved in human and mouse (Fig. S12A, using the following guides: GGTATTTGTACTCTAATGAA (guide 1), TGTACTCTAATGAAAGGAAC (guide 2), CTCACACAGGACAATCCGGCAGG (guide 3). For genotyping, we used primers (forward 1: 5-GAAATCGCATGTCACGCAGAAAT-3, reverse 2: 5-ATATCGGAACTGACATACTGAACG-3, reverse 2.2: 5- GTGAGCTTCGTGTTTGTGGT-3) to amplify a region around the transcription start site.

*Serpine3*^cbg18^ was generated by introducing a frame shifting deletion in coding exon 1, which presumably results in three early stop codons when the transcript is translated. The following guides were used: TCTTCTGCAACTCGGGGCCA (guide 4), TCTCTGTGAGCGTCTGGTAG (guide 5), AACACTCTGGTTCAGCTCGA (guide 6) (Fig. S12). We genotyped fish by amplifying a region around coding exon1 with the following primers: forward 3: 5- GGCATTGTTGAGATTCAGTAGTCA-3, reverse 4: 5-CAGTTTACTCCTACCATTGACATC-3.

### Histology

For hematoxylin/eosin stainings, fish were sacrificed and heads were fixed overnight at 4°C in 4% paraformaldehyde/ 0.1 M PBS and decalcified in 0.5 M EDTA in 0.1 M PBS for 3-4 days. Next, they were processed in a Paraffin-Infiltration-Processor (STP 420, Zeiss) according to the following program: ddH_2_0: 1×1′; 50% ethanol (EtOH) 1×5′; 70% EtOH 1×10′; 96% EtOH 1×25′; 96% EtOH 2×20′; 100% EtOH 2×20′; xylene 2×20′; paraffin 3×40′/60°C; paraffin 1×60′/60°C. The heads were embedded in paraffin using the Embedding Center EG1160 (Leica). Semi-thin sections (2 µm) were cut on an Ultracut microtome (Mikrom) and counterstained using hematoxylin/eosin (HE, Sigma).

### Microscopy, image processing and analysis

Imaging was performed using the ZEISS Axio Imager.Z1 provided by the CMCB Light Microscopy Facility. The images were processed in Fiji/ImageJ version 2.1.0 (110) (macroscopic eye images) and Adobe Illustrator. For macroscopic phenotyping, eyes were imaged with a Leica stereo microscope M165C. To parameterize the eye shape, the eye outline was first approximated by an oval and then manually corrected if necessary. Particle parameters of the final eye object were measured automatically in Fiji. The statistical analysis and visualization were conducted in R version 4.1.0 (2021-05-18) using the packages ggplot2 (111) and tseries (112). Comparing WT and KO individuals, we tested whether both genotypes have the same iris shape (estimated by iris solidity and circularity) using the Wilcoxon rank sum test after rejecting normality of the variables with a Jarque Bera Test. Both eyes of the same individual were treated as individual biological replicates, since we observed shape deviations often only in one eye of the same individual (Tab. S7).

### Animal licenses

All experiments in mouse and zebrafish were performed in accordance with the German animal welfare legislation. Protocols were approved by the Institutional Animal Welfare Officer (Tierschutzbeauftragter), and licensed by the regional Ethical Commission for Animal Experimentation (Landesdirektion Sachsen, Germany; license no. DD24-5131/354/11, DD24.1-5131/451/8, DD24-5131/346/11, DD24-5131/346/12).

### Data availability

All data needed to evaluate the conclusions in the paper are present in the paper and the Supplementary Materials. The phylogenetic tree used for the selection, the annotated protein alignment of mammalian SERPINE3, and the predicted structure of human SERPINE3 is available at https://genome.senckenberg.de//download/SERPINE3/.

## Acknowledgment

We thank the genomics community for sequencing and assembling the genomes and the UCSC genome browser group for providing software and genome annotations. Experimental work would not have been possible (or as pleasant) without supporting hosting labs, especially Nadine Vastenhouw, Elisabeth Knust and Wieland Huttner. We also thank Nadine Vastenhouw and her whole lab, Michael Heide and Mauricio Rocha as well as current and former members of the Hiller lab for helpful scientific discussion and comments on the manuscript. We thank the following facilities of MPI-CBG: Biomedical Services (especially fish unit), Cell technologies (Julia Jarrells), Sequencing and genotyping (Sylke Winkler), Light Microscopy, Scientific Computing, Computer Service Facilities as well as the Computer Service Facilities of MPI-PKS and the CMCB Histology (Susanne Weiche) and CMCB Light Microscopy Facility for their support. Work of HI and MH was supported by an exploration grant from the Boehringer Ingelheim Stiftung, the Max Planck Society and the LOEWE-Centre for Translational Biodiversity Genomics (TBG) funded by the Hessen State Ministry of Higher Education, Research and the Arts (HMWK). Work by JH, AM, SH and MB was supported by project grants of the German Research Foundation (Deutsche Forschungsgemeinschaft, project numbers BR 1746/3 and BR 1746/6) and an ERC advanced grant (Zf-BrainReg) to MB. This study was furthermore supported with a PhD scholarship from Studienstiftung des deutschen Volkes to JH.

## Abbreviations

KO: knockout
WT: wild type
RPE: retinal pigment epithelium
MG: Mueller glia
RT-qPCR: reverse transcription quantitative PCR
RNA-Seq: RNA-Sequencing
SNP: single nucleotide polymorphism
ISH: *In situ* hybridization
FDR: false discovery rate

## Supplementary Information Text

### Mammalian SERPINE3 have features of inhibitory, secreted SERPINs

Mammalian SERPINE3 carry an N-terminal signal peptide (Fig. S8) that is predicted to lead to their secretion into the extracellular space in 96% of all analyzed intact and complete SERPINE3. SignalP 5.0 did not predict the presence of a signal peptide for four species: fat dormouse, black flying fox and puma.

Sequence analysis of intact and complete SERPINE3s revealed that two key features of inhibitory serpins are well conserved among placental mammals. First, the substrate determining residues P4-P4’ are conserved (positions 366-372 in human SERPINE3), whereby P1 denotes the substrate binding scissile bond, position 369 (Fig. S9) (1). This position is occupied by an arginine in SERPINE3 as is the case in the inhibitory SERPINE1 and SERPINE2 proteins. Second, the close-by hinge region (positions 355-361) is mostly occupied by small amino acids without prolines. This may allow the insertion of the hinge region into the A beta sheet, a key feature of serpins’ inhibitory mechanism (2).

Furthermore, AlphaFold2 (3) predicted a three-dimensional structure of the human SERPINE3 that is very similar to other native serpins (mean RMSD to native structures 1.6 A, Tab. S9) and adopts the native fold of serpins with an exposed, disordered reactive core loop for substrate binding that does not seem to adopt an alpha-helical conformation as in the non-inhibitory ovalbumin (4). Taken together, this suggests that SERPINE3 functions as a secreted serine protease inhibitor.

---

**Fig. S1.**
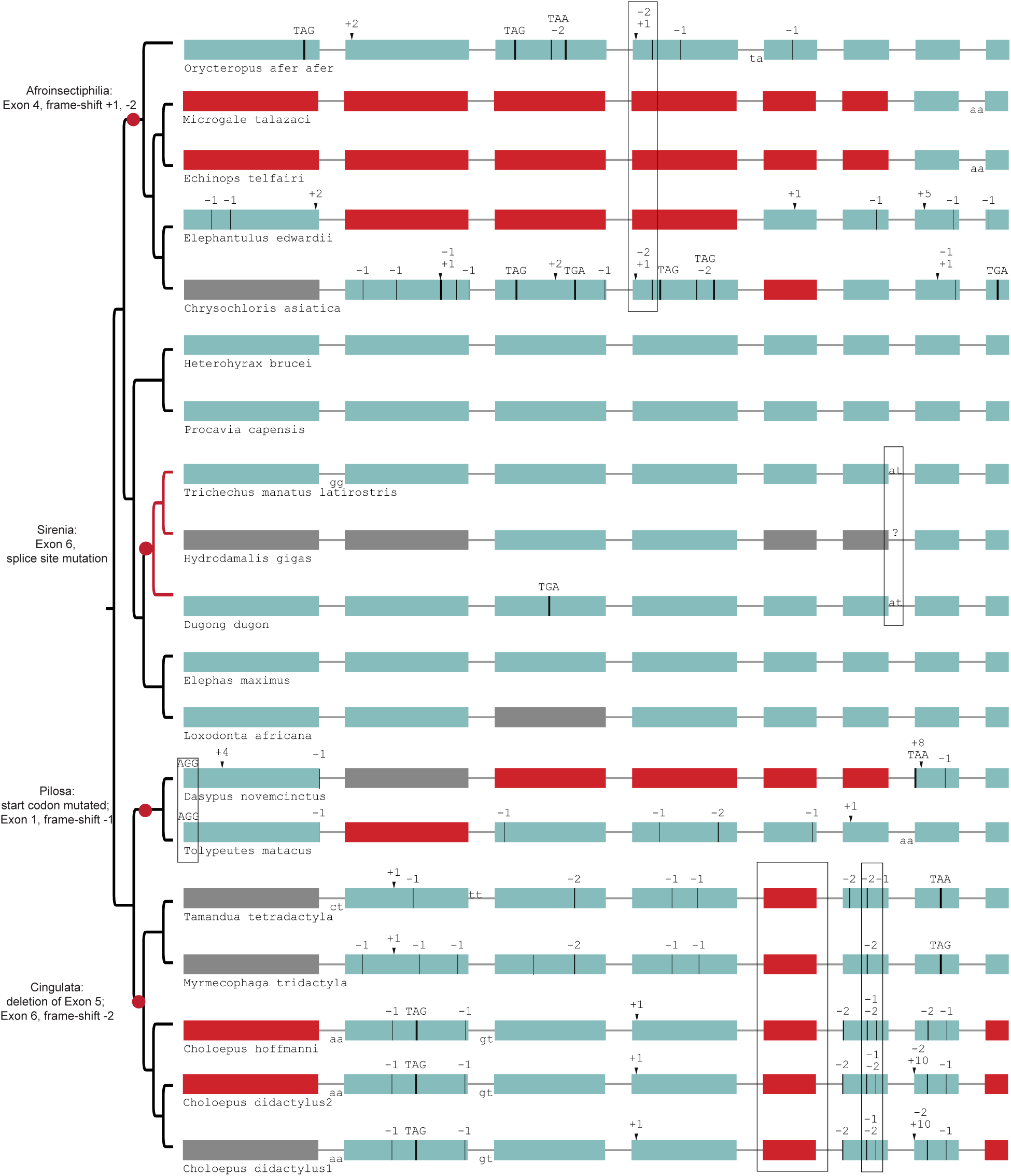
Gene-inactivating mutations support four independent losses of *SERPINE3* in Afrotheria and Xenarthra (red dots). Those shared mutations are marked by boxes, which indicate the loss of *SERPINE3* in the common ancestor of related species according to parsimony. For tenrecs (*Microgale talazaci*, *Echinops telfairi*) and elephant shrew (*Elephantulus edwardii*), the putative region of exon 4 was likely deleted after the shared frame-shifting mutations. *SERPINE3* in Sirenia (branches marked in red) evolve under relaxed selection. Coloring and legend as in Fig. 2 in the main text. Gray exons denote missing information.

**Fig. S2.**
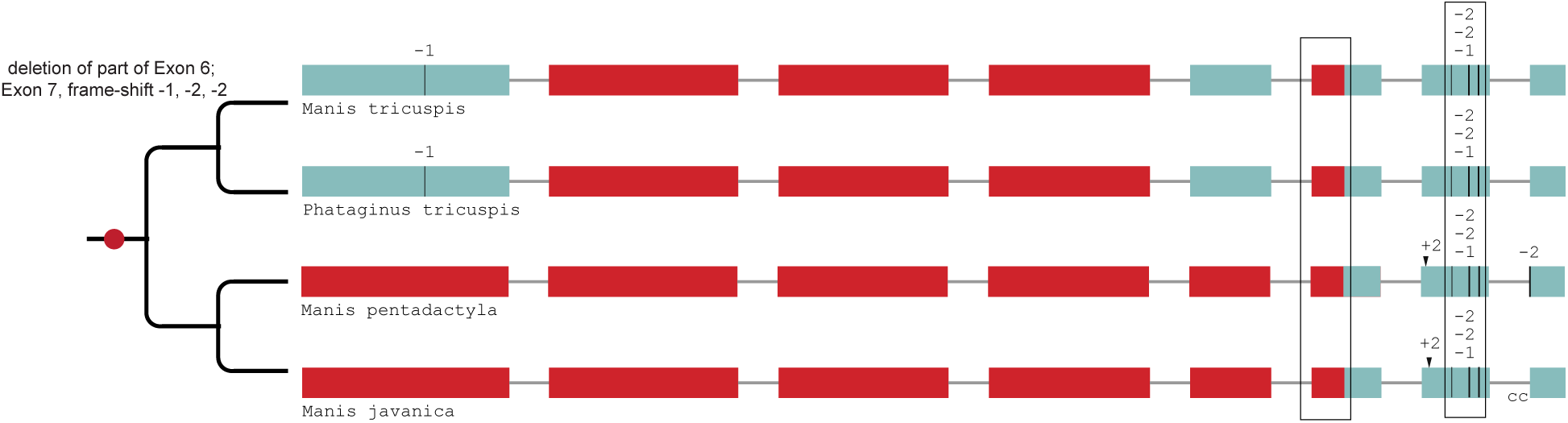
Gene-inactivating mutations support a single loss of *SERPINE3* in Pholidota. A partial deletion of coding exon 6 and three frame-shifting deletions in exon 7 are shared among all four species, indicating a single loss of *SERPINE3* in the Pholidota lineage. Coloring and legend as in Fig. 2 in the main text.

**Fig. S3.**
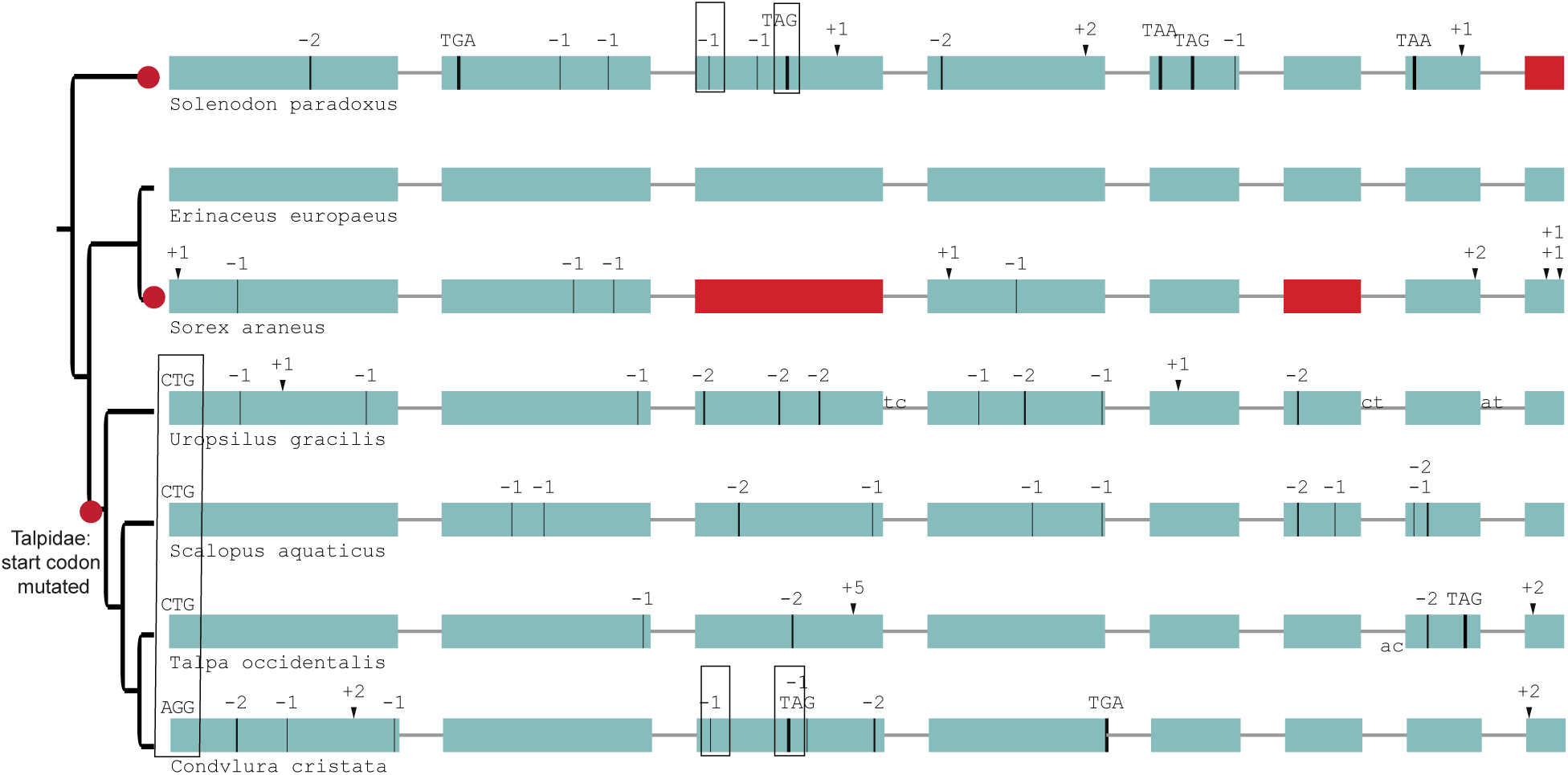
Gene-inactivating mutations support three independent losses of *SERPINE3* in Eulipotyphla (red dots). The start codon is not intact in the four moles (a). Star-nosed mole (*Condylura cristata*) and Hispaniolan solenodon (*Solenodon paradoxus*) share two inactivating mutations in exon 3, although they are phylogenetically distant (b, c). Coloring and legend as in Fig. 2 in the main text.

**Fig. S4.**
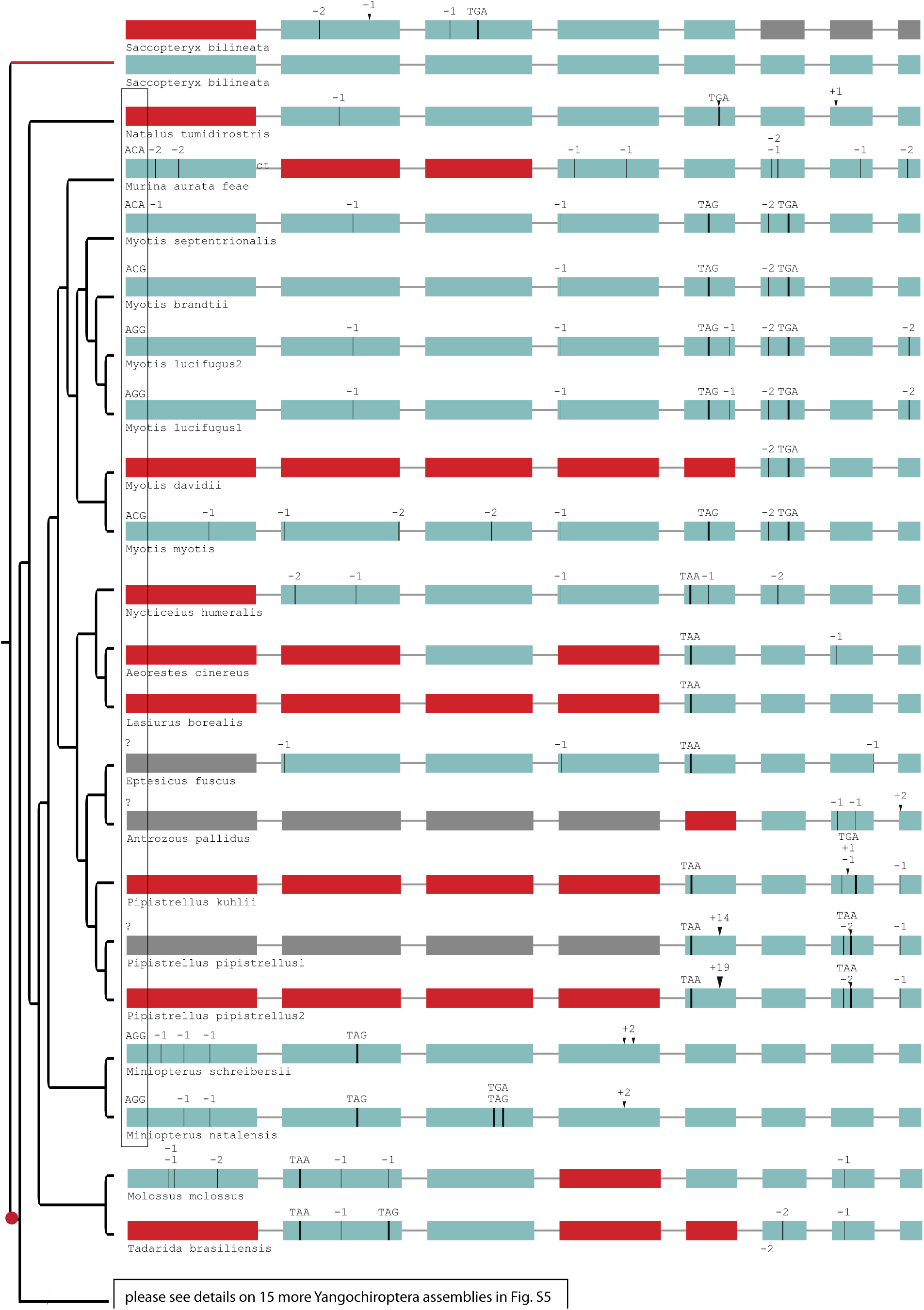
Gene-inactivating mutations support one loss of *SERPINE3* in Yangochiroptera. The ancestral start codon is mutated in most Yangochiroptera (box). The most parsimonious explanation is a shared start codon mutation (red dot) in the ancestral lineage after split from the sac-winged bat (*Saccopteryx bilineata*). A back mutation to the regular start codon ATG likely occurred in *Molossus molossus* and *Artibeus jamaicensis*. Please see Fig. S5 for gene-inactivating mutations of 15 more Yangochiroptera assemblies. Sac-winged bat has an intact copy of the gene that evolves under relaxed selection (red branch) in addition to a second copy of *SERPINE3,* which was independently inactivated. Coloring and legend as in Fig. 2 in the main text. Gray exons denote missing information.

**Fig. S5.**
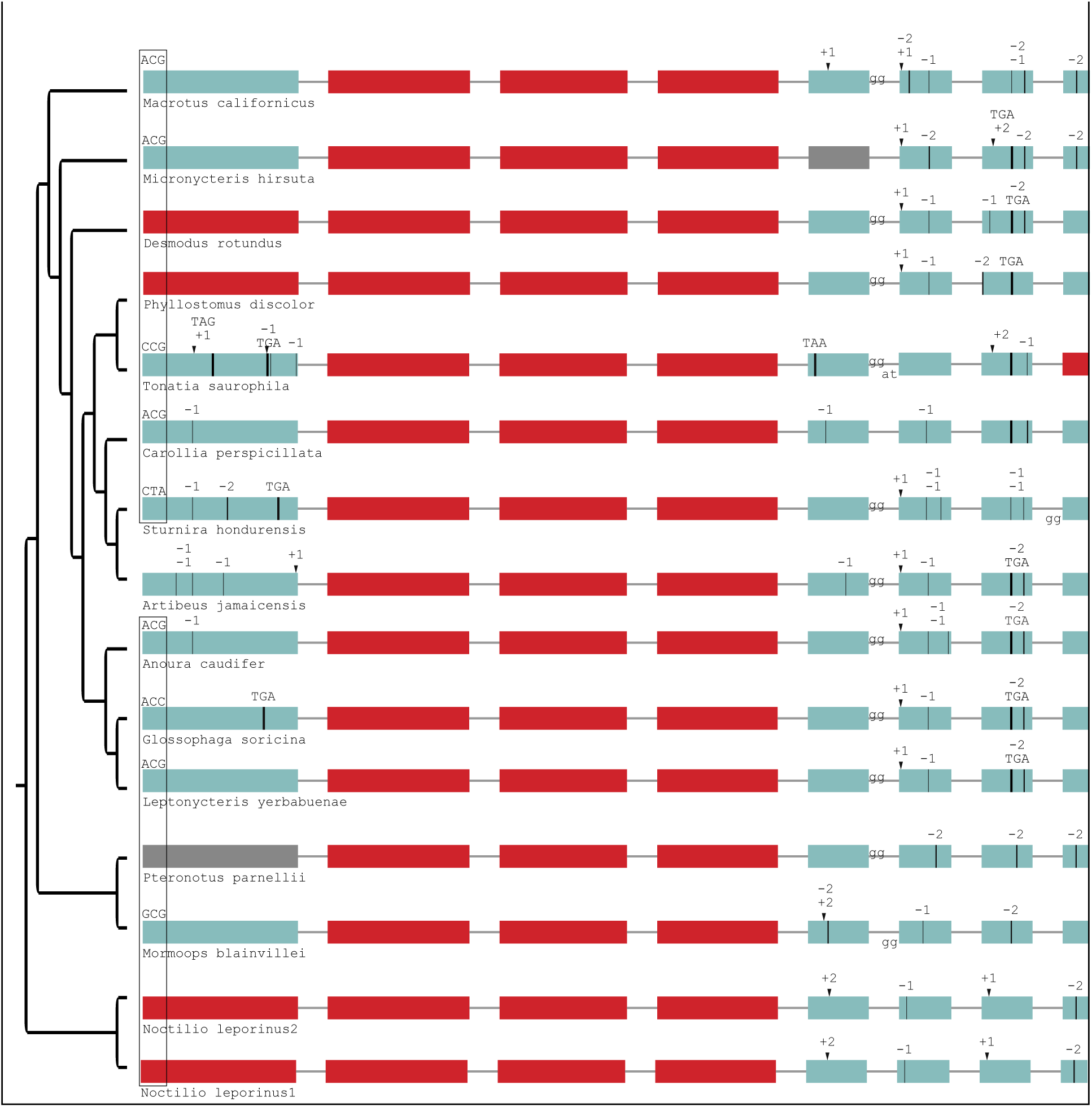
Gene-inactivating mutations in more Yangochiroptera. This is an extension to Fig. S4. The start codon mutation shown is shared with most other Yangochiroptera (Fig. S4). Coloring and legend as in Fig. 2 in the main text. Gray exons denote missing information.

**Fig. S6.**
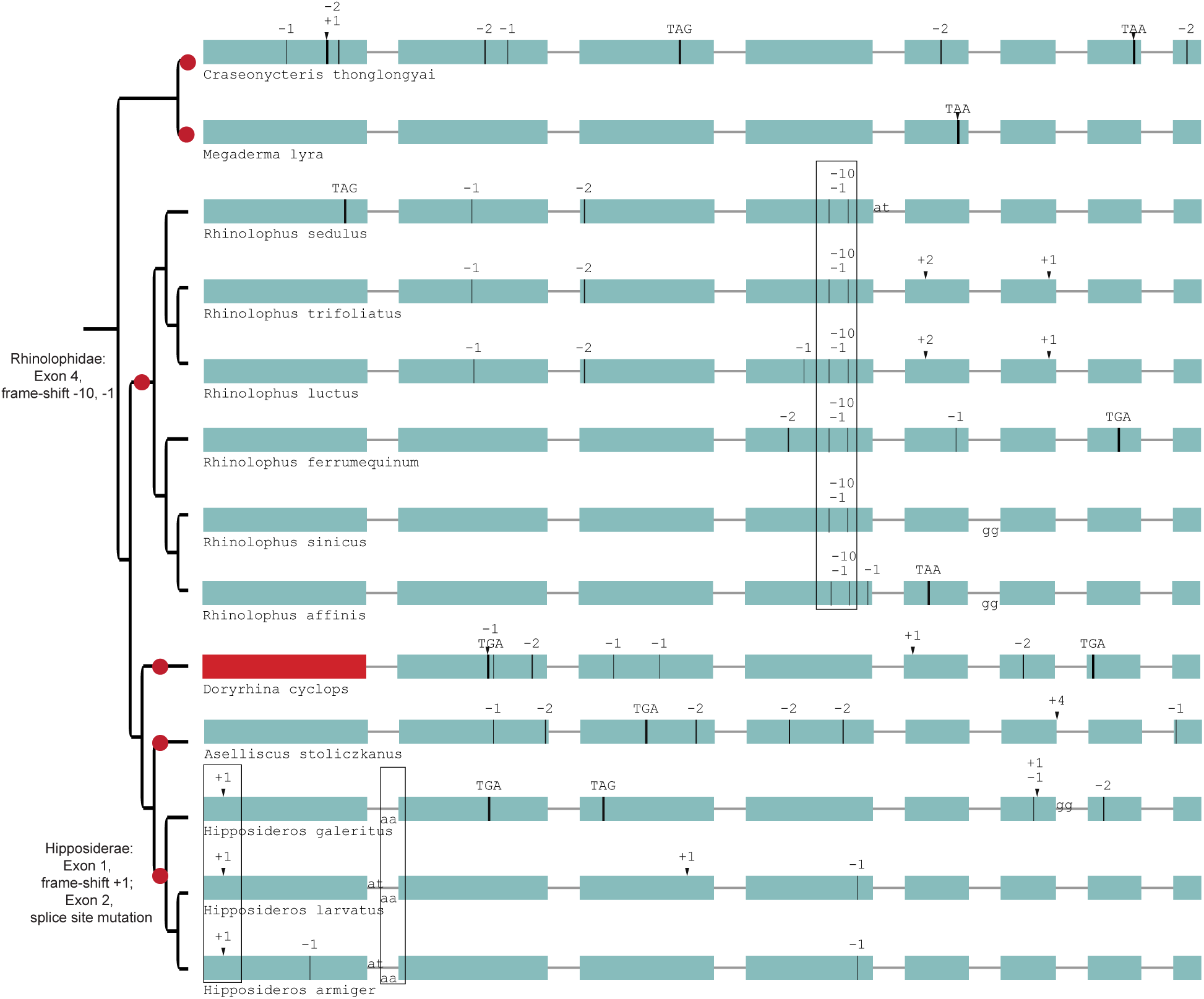
Gene-inactivating mutations support six independent losses of *SERPINE3* in Yingochiroptera. Six independent inactivation events (red dots) occurred in *SERINE3*’s coding sequence in Yingochiroptera. All Rhinolophidae share frame-shifting mutations in exon 4 (boxed), while Hipposideros share a frame-shifting insertion in exon 1 and a splice site mutation at exon 2 (boxed). Coloring and legend as in Fig. 2 in the main text.

**Fig. S7.**
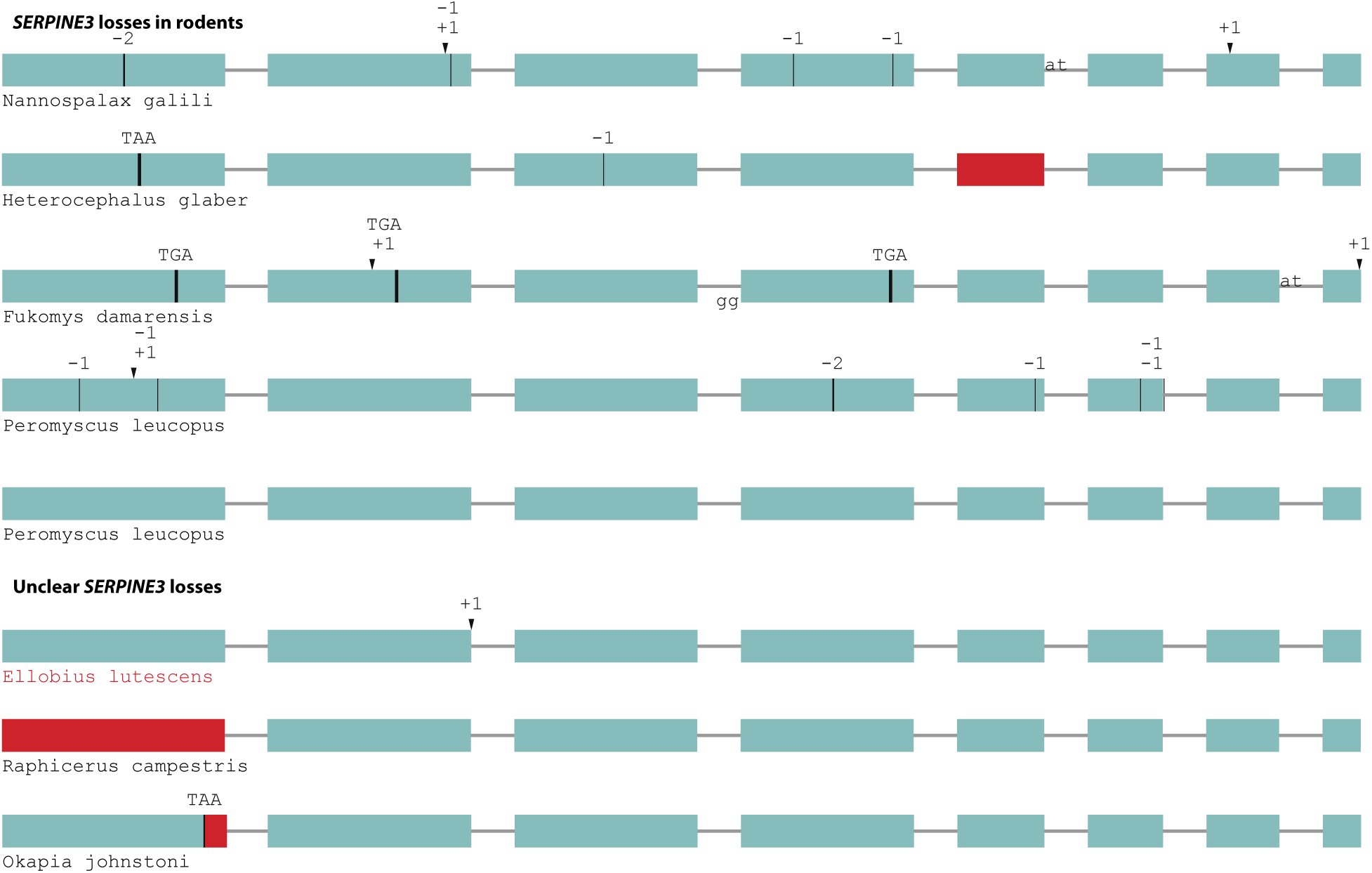
Independent gene-inactivating mutations in other mammals. A test for shifts in selective pressure revealed that *SERPINE3* evolves under relaxed selection in the subterranean mole vole (*Ellobius lutescens*, marked in red), but not in the other cases of gene-inactivating mutations with unclear consequences (steenbok, okapi). A species-specific duplication with inactivation of one *SERPINE3* copy occurred in white-footed mouse (*Peromyscus leucopus*). Coloring and legend as in Fig. 2 in the main text.

**Fig. S8.**
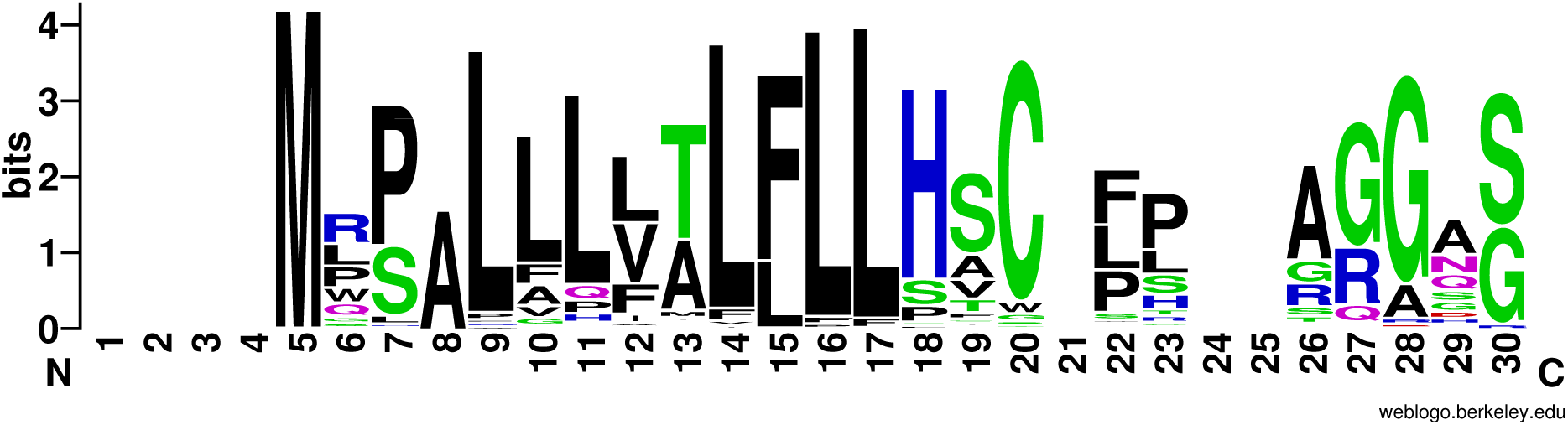
Conservation of the signal peptide in mammalian SERPINE3. The putative N-terminal signal peptide contains many hydrophobic residues in intact mammalian SERPINE3 and is predicted to guide secretion into extracellular space. We show the first 30 alignment columns. The sequence logo was generated with Weblogo (5).

**Fig. S9.**
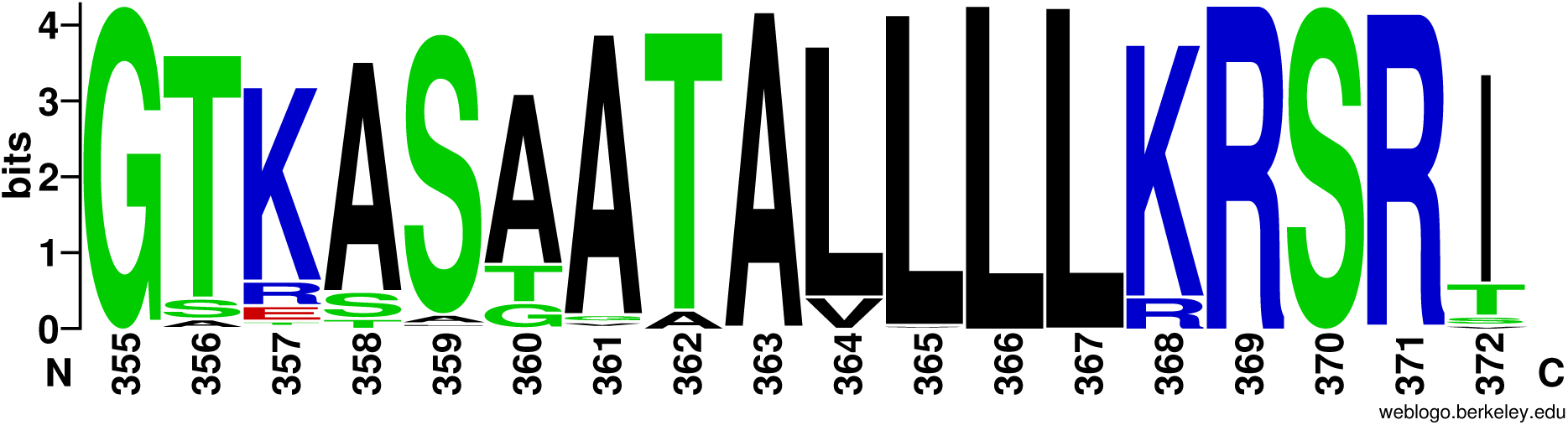
The hinge region and reactive core loop are conserved in intact mammalian SERPINE3. The numbering is in reference to human SERPINE3, where R369 likely is the scissile bond (P1) within the reactive core loop (positions 366-372, P4-P4’). The hinge region is located at positions 355-361. The logo was generated with Weblogo (5).

**Fig. S10.**
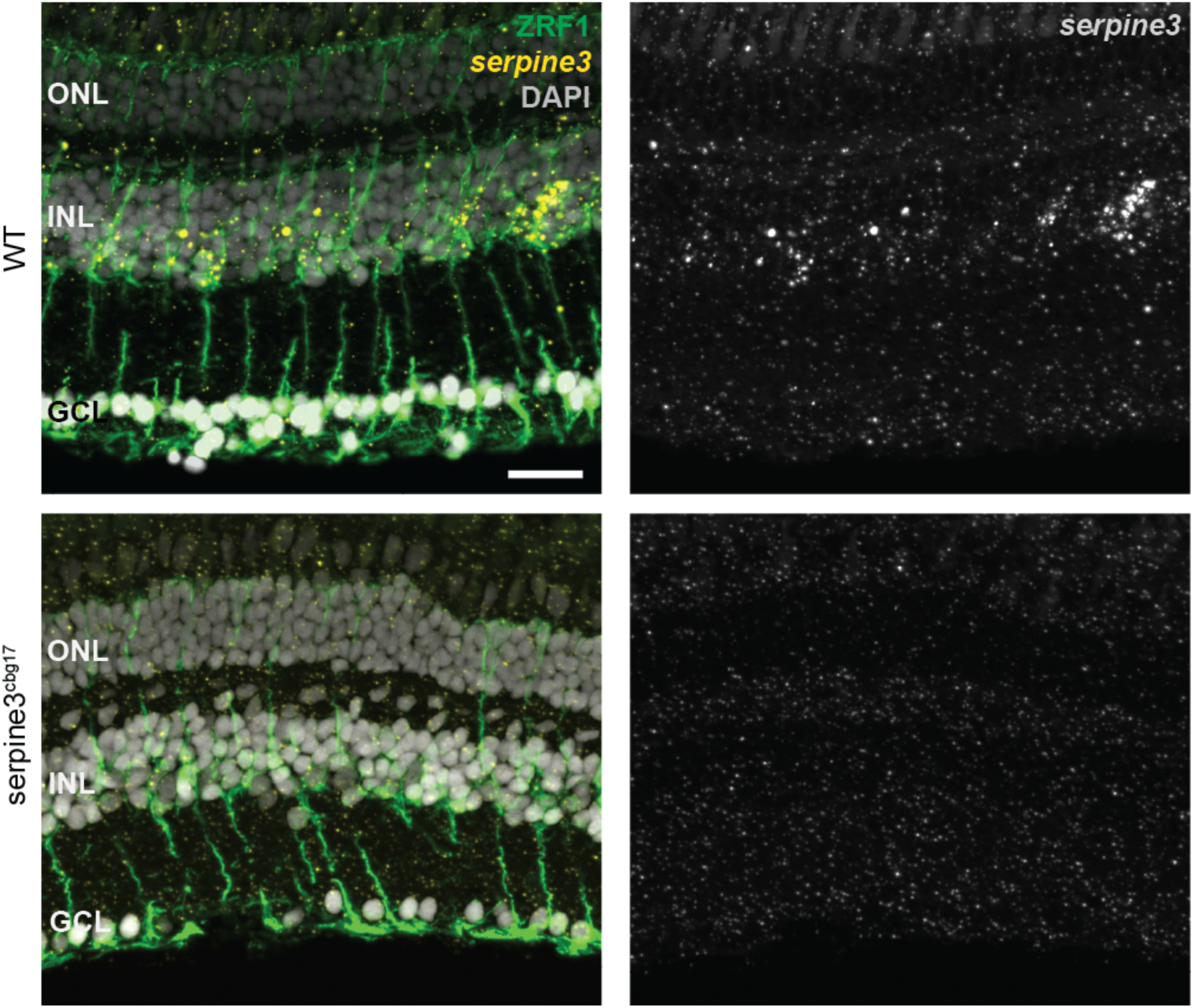
*Serpine3* expression co-localizes with glial fibrillary acidic protein in the retina of wild type fish. The fluorescence *in situ* signal for zebrafish *serpine3* is specific for wild type (WT, yellow), where it is in proximity to staining of the ZRF1-antibody (green). This indicates expression of *serpine3* by Mueller glia cells. *Serpine3* signal is not present in homozygous *serpine3*^cbg17^ siblings. In the overlay, nuclei (DAPI) are shown in white. ONL – outer nuclear layer, INL – inner nuclear layer, GCL – ganglion cell layer.

**Fig. S11.**
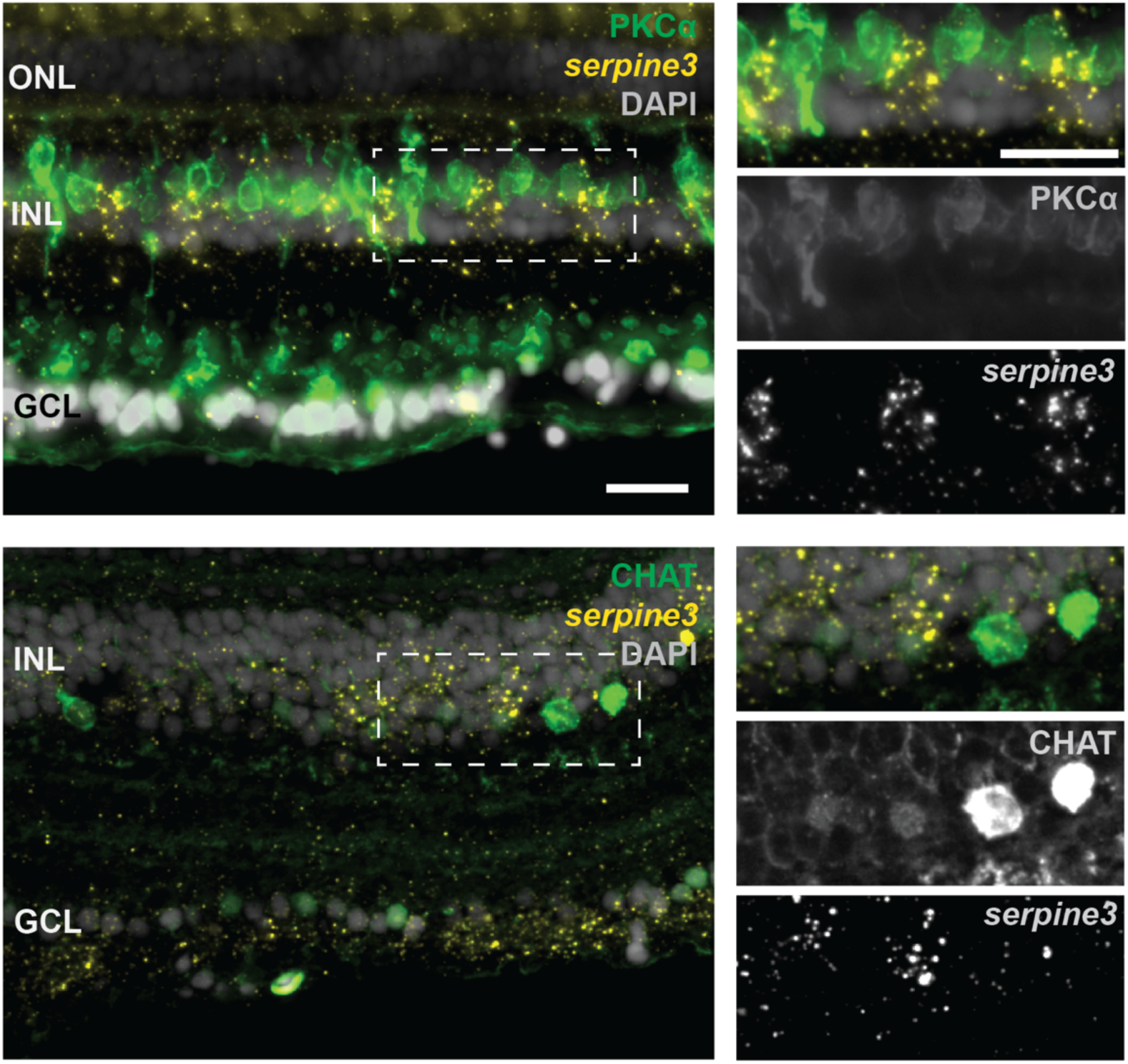
*Serpine3* expression does not co-localize with markers for bipolar or amacrine cells in the zebrafish retina. Fluorescence *in situ* hybridization of *serpine3* mRNA (yellow) does not show co-localization with anti-protein kinase C alpha (PKCa encoded by *prkca*, green) or choline O-acetyltransferase a (CHAT, encoded by *chata*, green) proteins. In the overlay, nuclei (DAPI) are shown in white. ONL – outer nuclear layer, INL – inner nuclear layer, GCL – ganglion cell layer. Scale bar overview = 20 µm, details = 20 µm.

**Fig. S12.**
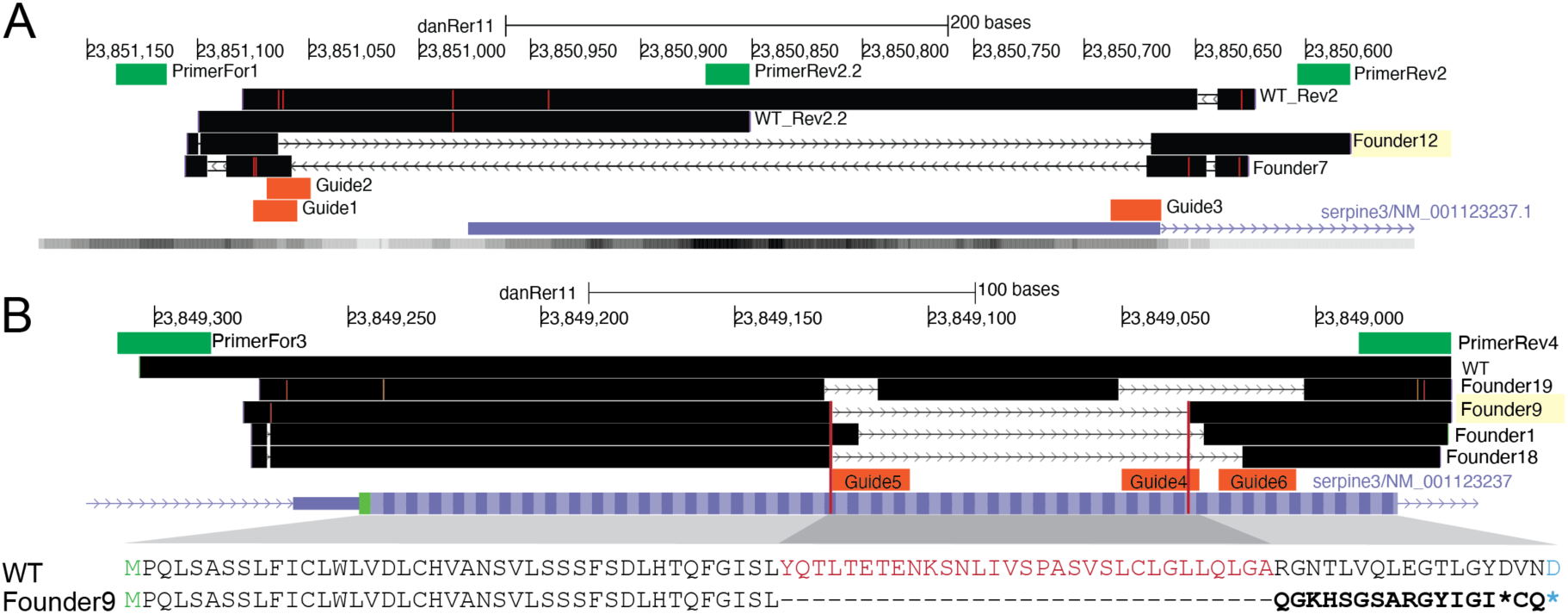
Knockout of *serpine3*^cbg17^ and *serpine3*^cbg18^ alleles in zebrafish by CRISPR-Cas9 is confirmed by sequencing results. Sequenced reads are mapped against the danRer11 zebrafish genome assembly with Blat and are visualized in the UCSC genome browser (black) together with the *serpine3* RefSeq annotation (blue). (A) PCRs with primers For1 and Rev2 or For1 and Rev2.2 (green) amplify the expected regions around the transcription start site on chromosome 9 in wild type (WT_Rev2, WT_Rev2.2). Injection of CRISPR guides 1, 2 and 3 (orange) results in a deletion of about 400 bp for founder individuals 7 and 12 in *serpine3*^cbg17^. Offspring of founder 12 (394 bp deletion) was raised and further crossed. The location of a single *serpine3* transcription start site is supported by annotation and activating histone marks H3K4me3 within the respective region (lower gray bar with darkness of Color correlating with signal intensity). (B) PCR with primers For3 and Rev4 (green) amplifies a region around coding exon1 of *serpine3*. Injection of CRISPR guides 4, 5 and 6 (orange) results in a deletion of about 100 bp for founders 1, 9, 18 and 19 in *serpine3*^cbg18^. For founder 9, this 92 bp deletion induces a frame-shift in the reading frame (bold) with three early stop codons (two shown as *, third stop located in coding exon 2). The deletion is equivalent to deletion of 31 amino acids (red) and +1 nt insertion. The amino acid encoded by a split codon is shown in blue.

**Fig. S13.**
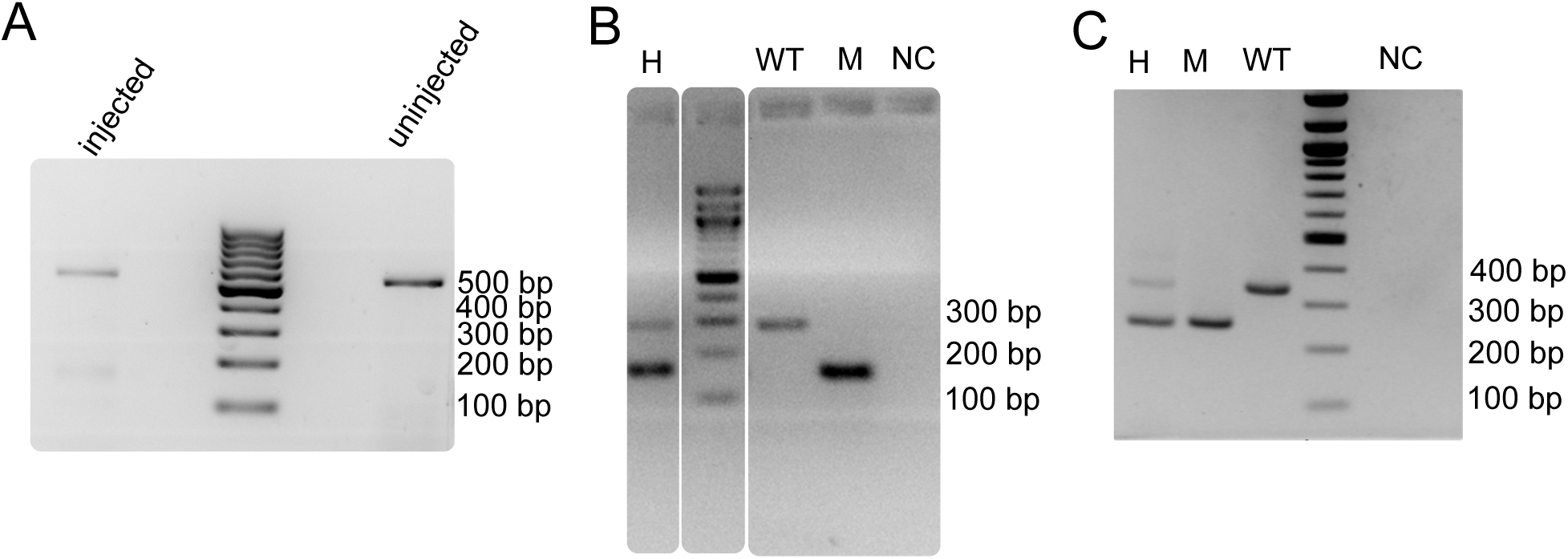
PCRs confirm expected CRISPR-Cas9 induced deletions in *serpine3*^cbg17^ and *serpine3*^cbg18^ fish. (A) Genotyping of the *serpine3*^cbg17^ with two primers (F0 generation). Injection of CRISPR guides into the one-cell embryo leads to mosaic embryos (several pooled at 72 hpf) with a strong wild type (WT) band at about 557 bp and a weak mutant band at about 161 bp, while uninjected embryos just have a single WT band (557 bp). (B) Genotyping of *serpine3*^cbg17^ with a mix of three primers (F2 generation). Heterozygous animals (H) have two bands, a WT (expected height: 286 bp) and mutant band (expected height: 161 bp), while homozygous mutants (M) and homozygous WT animals show a single band. (C) Genotyping of *serpine3*^cbg18^ with two primers. Heterozygous animals (H) have two bands, a WT (expected height: 342 bp) and mutant band (expected height: 250 bp), while homozygous mutants (M) and homozygous WT animals show a single band. NC – negative control (water).

**Fig. S14.**
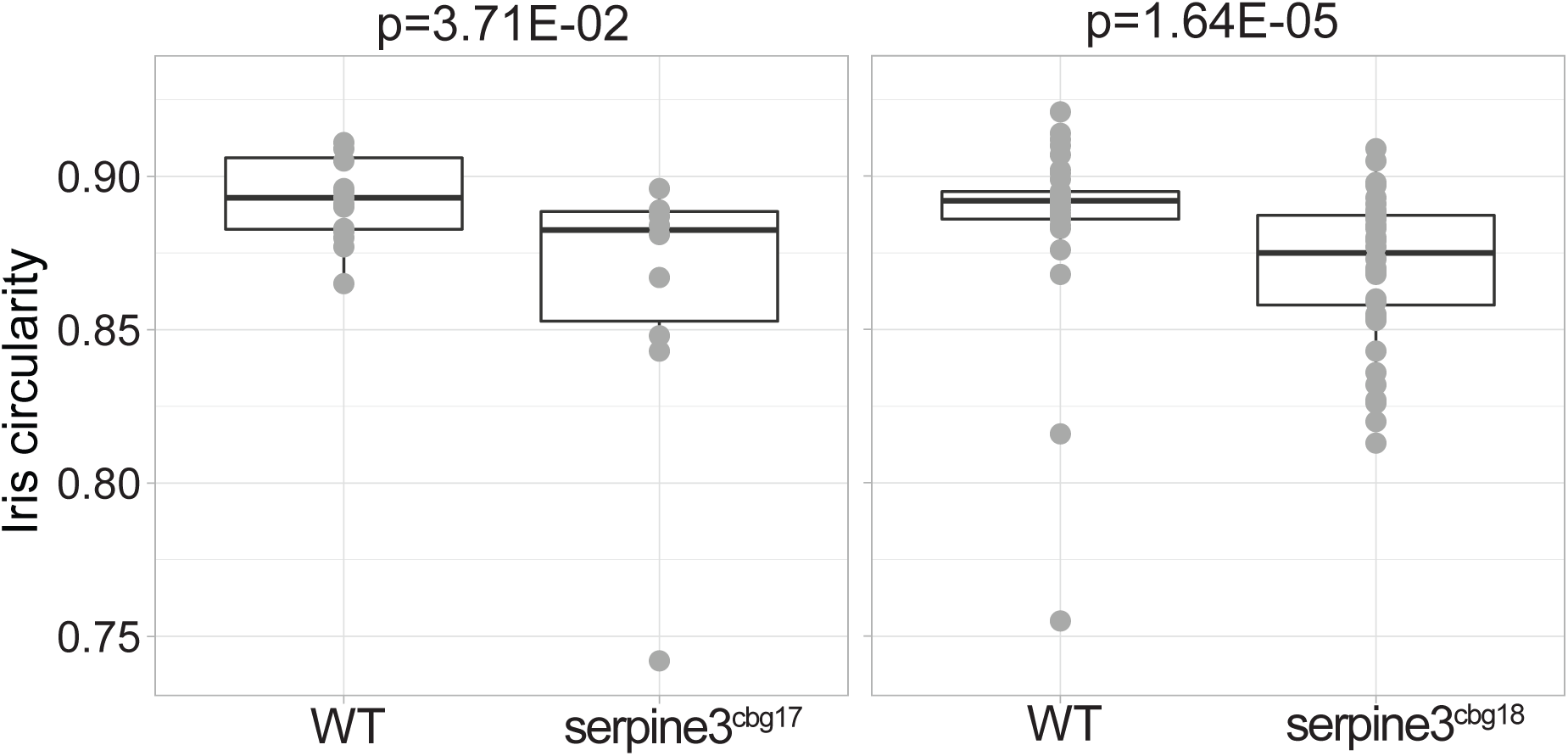
Iris circularity differs between *serpine3*^cbg17^ and *serpine3*^cbg18^ fish and their respective wild type siblings (WT). Iris circularity is another descriptor for the deviation of eye shape and is defined as 4*π* × [Area]/[Perimeter]^2^. A value of 1 indicates a perfect circle. Box plots show that iris circularity significantly differs between WT (n=14) and *serpine3*^cbg17^ eyes (n=10). The same holds for the comparison of WT (n=40) and *serpine3*^cbg18^ eyes (n=40). A Wilcoxon Rank sum test was used.

**Fig. S15.**
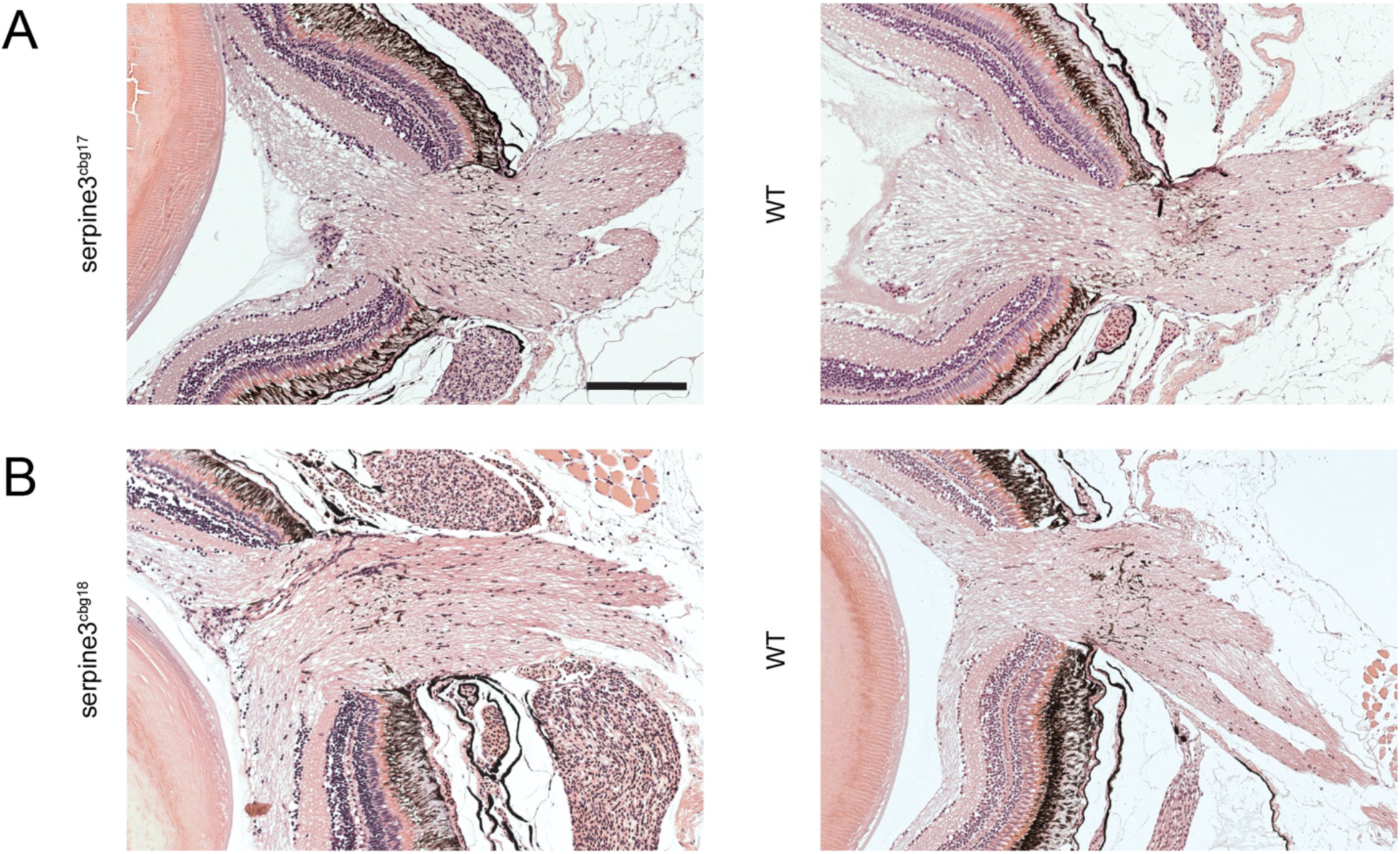
The optic nerve is intact in *serpine3*^cbg17^ and *serpine3*^cbg18^ fish as shown in a hematoxylin/eosin histology staining. No difference in optic nerve morphology is observed between *serpine3*^cbg17^ (A) and *serpine3*^cbg18^ fish (B) and their respective wild type (WT) siblings. Scale bar = 200 µm.

**Fig. S16.**
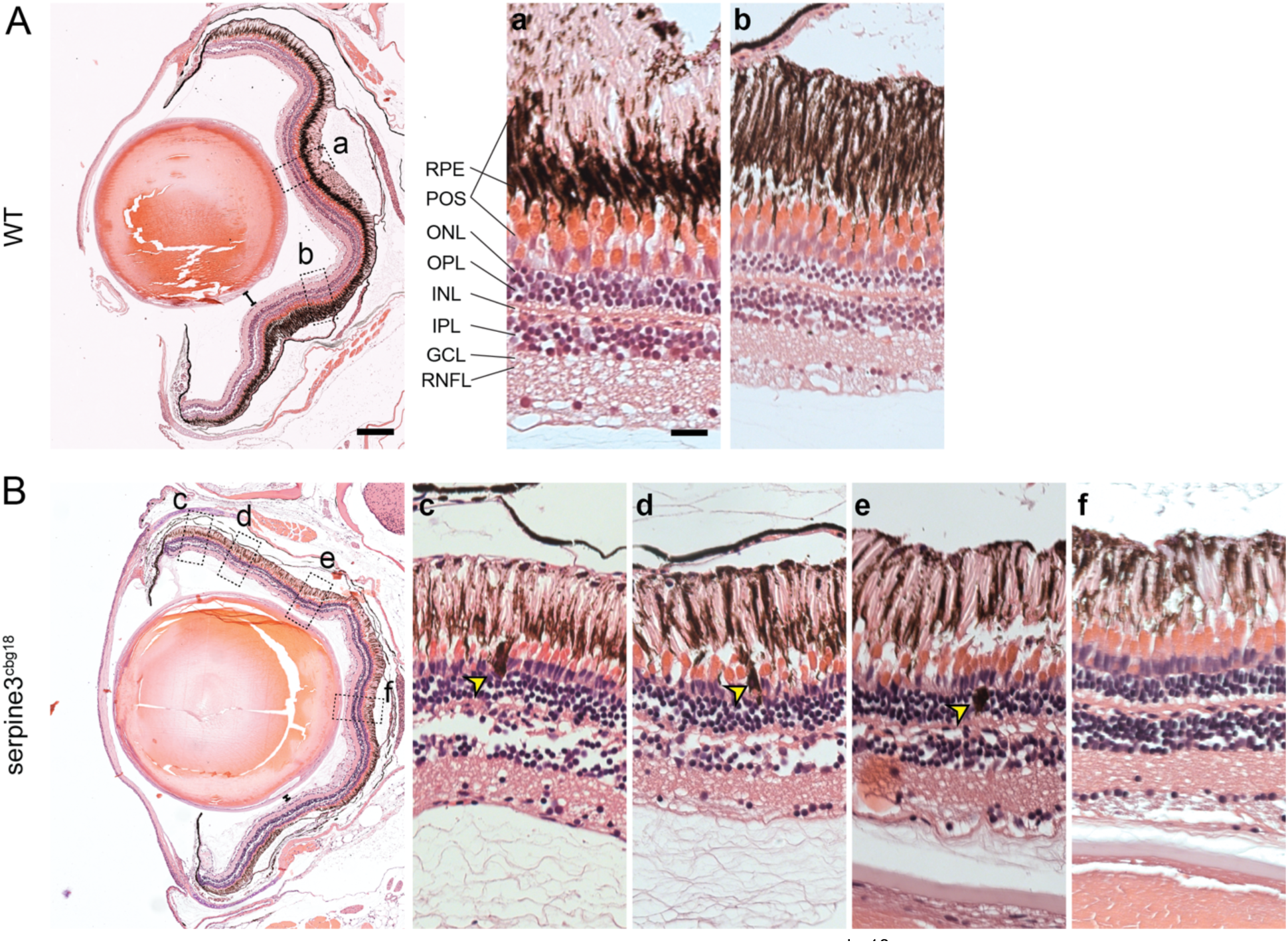
Hematoxylin/eosin histology staining of *serpine3*^cbg18^ eyes (14 months) reveals histological differences in comparison to their wild type (WT) siblings (dorsal top, ventral bottom). We show details of representative overview images for one eye of each genotype on the right. In comparison to WT, distance between lens and retina of *serpine3*^cbg18^ fish is reduced (distance bars). In the *serpine3*^cbg18^ eye (B), all retinal layers are present and distinguishable, although they are not as tightly packed and clearly separated as in the WT (c-f). Moreover, we observed displaced pigmented cells located mainly in the photoreceptor outer segment and the outer nuclear layer (yellow arrows, c-e). Furthermore, the photoreceptor outer segment and RPE layer are not clearly separated in the *serpine3*^cbg18^ retina. Scale bar in the overviews represents 200 µm and in the magnifications 20 µm.

